# Static2Dynamic: Reconstructing videos of unobservable cellular, developmental, and disease processes

**DOI:** 10.64898/2026.05.18.725860

**Authors:** Thomas Boyer, Elaine del Nery, Nathalie Spassky, Auguste Genovesio

## Abstract

A fundamental limitation in biology is that many of its most important processes unfold as visual dynamics that cannot be directly observed. Development, tissue remodeling, and disease progression often occur deep in living organisms, over extended timescales, and at cellular resolution beyond the reach of current live imaging technologies. As a result, much of biology remains accessible only through static snapshots, while the underlying phenotypic trajectories and visual transformations remain hidden.

Here, we introduce Static2Dynamic, a general framework to reconstruct unseen biological dynamics from sets of cross-sectional image data. Starting from time-unpaired static samples, Static2Dynamic first recovers a continuous pseudotime for individual images in a time-discriminative deep representation space, then learns a generative model of images conditionally to the underlying process, and finally reconstructs temporally coherent videos initialized from real samples. This makes it possible to infer past and future visual states of a static image and to simulate complete trajectories of cellular, developmental, and disease processes that were never directly recorded.

We quantitatively validate *Static2Dynamic* on two large-scale experimental microscopy video datasets generated specifically for benchmarking, enabling direct comparison of inferred pseudotime trajectories and reconstructed videos against ground-truth biological dynamics. We further show that the framework generalizes across biological scales, organisms, and imaging modalities, including processes inaccessible to continuous live observation. More broadly, *Static2Dynamic* establishes the foundations of *pseudotime microscopy*, a new paradigm for reconstructing the visual and temporal dynamics of biological processes directly from static imaging data, thereby expanding the observable space of living systems beyond current experimental limits.

## Introduction

As an experimental science, biology heavily relies on observation. Technological progress over the centuries has continuously augmented our ability to observe and describe phenotypic traits across species. In the past few decades, revolutionary methods such as super resolution imaging, high throughput sequencing or mass spectrometry, to name a few, have enabled unprecedented quantification of phenotypic variations at cellular and molecular scales, leading to transformative discoveries (Betzig et al. 2006; Bentley et al. 2008; Aebersold and Mann 2003).

However, important limitations remain in our ability to observe biological systems, particularly when it comes to continuously monitoring cells, tissues or organs in their natural environment –in vivo. While time-lapse microscopy of cell culture and certain organism models have been made possible for decades, in vivo spatial or visual observation of cellular and long lasting processes in most living tissues remains largely out of reach. This is the case for instance with the visualization of morphological transformations occurring during in utero development, or disease progression in deep tissues. On one hand, continuous visual observation of cells is largely impossible in vivo because current fluorescent microscopy techniques cannot resolve individual cells in non-transparent tissues deeper than about half a millimeter (Razansky et al. 2009). On the other hand, monitoring most long lasting developmental processes or disease progression –which can take weeks, months or even years– is unfeasible, as existing observation methods are often highly invasive, if not destructive. Even in the rare cases where non-destructive methods exist (Horton et al. 2013; Streich et al. 2021; Barretto et al. 2011; Wang and Hu 2012), they come with trade-offs such as reduced spatial or temporal resolution and limited visual coverage. Altogether, long-term or cellular-scale dynamics in deep tissues remain largely inaccessible. As a result, we are effectively blind to the spatial and visual aspects of most developmental and disease processes, which hinders our understanding of fundamental biology and limits progress in early diagnosis and prognosis.

While visual continuous observation of a process remains challenging, sets of ordered cross-sectional image data –that is: sets of samples collected from different individuals and acquired at discrete time points– can often be obtained for instance through a destructive measurement process over developmental days or as roughly annotated stages of a disease progression. Over the past decade, single cell sequencing of such static samples were used to reconstruct continuous transcriptomic variations occurring during differentiation processes (Cao et al. 2019; Lopez et al. 2018; Gayoso et al. 2024; Klein et al. 2025). Transcriptomic pseudotime provides unique access to complex molecular dynamic processes. For instance, it has been instrumental in uncovering receptor transformations during olfactory neurogenesis (Hanchate et al. 2015). These approaches benefit from the fact that gene expression counts are vectors that are meaningful cell profiles by construction: a short distance between two cell profiles directly reflects close gene expression. This enables fitting a curve or a graph directly within this representation but also directly interpreting the trajectory in terms of individual gene expression changes. By contrast, although collecting a set of cross-sectional image data is possible in many studies, pseudotime inference and individual path reconstruction in this image context faces significant challenges due to the high variability of image data and the lack of a robust time-discriminative representation at the pixel level. Moreover, naive interpolation techniques cannot be applied directly to images because, contrary to gene expression, a large distance between two images in pixel space does not imply that their associated phenotypes are different. Therefore, no generalizable approaches to recover pseudotime sequences from still images beyond those dedicated to specific analyses, such as the cell cycle (Rappez et al. 2020), have been attempted. More importantly, to our knowledge, reconstructing a video of the biological process from these static snapshots has never been achieved.

In this work, we introduce Static2Dynamic, a method in three stages that offers a robust and generalizable solution to this problem. Assuming we have only access to a small set of ordered cross-sectional image data, Static2Dynamic first estimates the continuous pseudotime of individual snapshot images, then learns a rich time-conditioned distribution of images from static time-unpaired samples, and finally generates temporally coherent videos from true starting samples, simulating their unseen dynamic for further visual observation. The pseudotime inference stage relies on a deterministic approach that projects all images on a cubic spline in a time-discriminative deep image representation space. The time-conditioned distribution learning and video generation steps rely on a diffusion model continuously conditioned on the pseudotime obtained in the first step. The whole method makes possible the reconstruction of the past and future video of the underlying process from any static image. We extensively validate this approach on two large experimental *video* microscopy datasets produced for evaluation purposes, where the pseudotime inference and the videos obtained from this method can actually be quantitatively compared to their actual ground truth video counterpart. We then illustrate its broad applicability at various biological scales, organisms and image modalities where ground truths are not accessible.

## Results

### Static2Dynamic: a method to generate videos from unpaired, static images

To recover the continuous nature of an underlying biological process from a dataset of cross-sectional image data, we first derived an approach to estimate, for each image, a continuous pseudotime value between 0 –the beginning of the full process– and 1 –its end. Pseudotime estimation itself comprises 3 stages (**Figure 1**A and Methods). Briefly, images are first encoded using DINOv2, an effective foundational model trained in a self-supervised way (Oquab et al. 2023). While this stage provides a highly discriminant representation, PCA and UMAP transforms show that it does not especially favor time variation over other features for most datasets (**Supp. Fig. 1**). To emphasize and combine the most time-varying discriminative features, we further compute a Linear Discriminant Analysis (LDA) from these encodings using cross section labels (say acquisition times) as classes. Such a transformation already enables us to see a clear general time evolution on most datasets (**Supp. Fig. 2**). Finally, in order to materialize it, a spline is fit to the LDA class centroids, yielding a smooth continuous pseudo-time curve. Projecting all samples onto this curve in the time-discriminative LDA space enables robust individual pseudotime estimation for each image of a dataset (**Figure 1**A and **Supp. Fig. 3**). This deterministic first stage of Static2Dynamic allows fine-grain reordering of each individual image along the underlying process, offering a generalizable and straightforward approach for pseudotime estimation.

**Figure 1:**
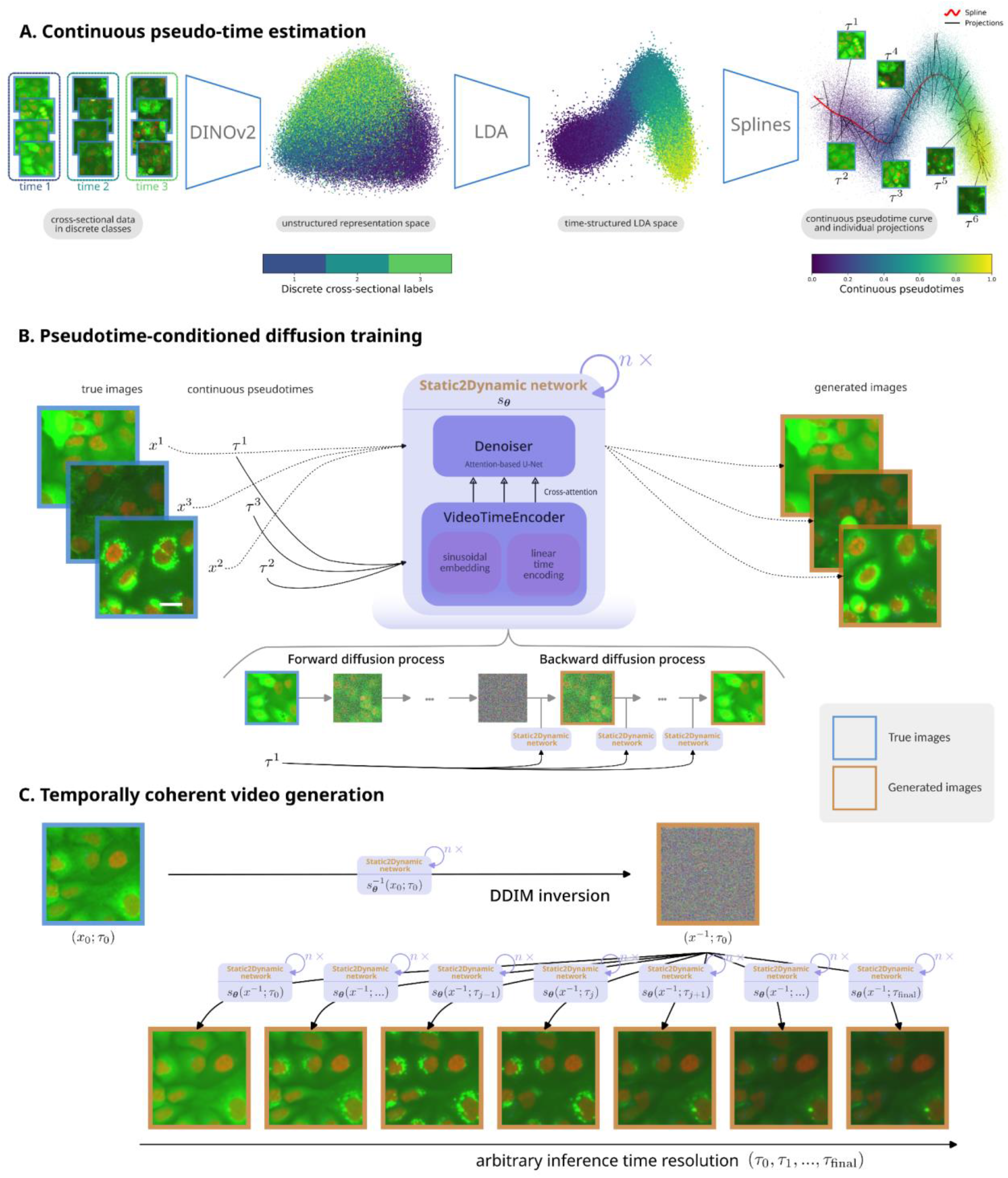
Description of our method. Static2Dynamic comprises 3 stages: A - continuous pseudotime estimation: DINOv2 encoding of images (PCA is shown here) is followed by an LDA transform, spline fitting on class centroids, and projection of the dataset on this spline to infer a pseudotime scalar from each image. B - a denoising diffusion image generator model is trained conditionally to the estimated pseudotime. The scale bar is 20 µm. C - at inference time, an image from the dataset is inverted to a gaussian latent using DDIM inversion and its estimated pseudotime, then an image sequence (aka a movie) can be obtained by conditioning the diffusion model with an arbitrary sequence of pseudotime values.

**Figure 2:**
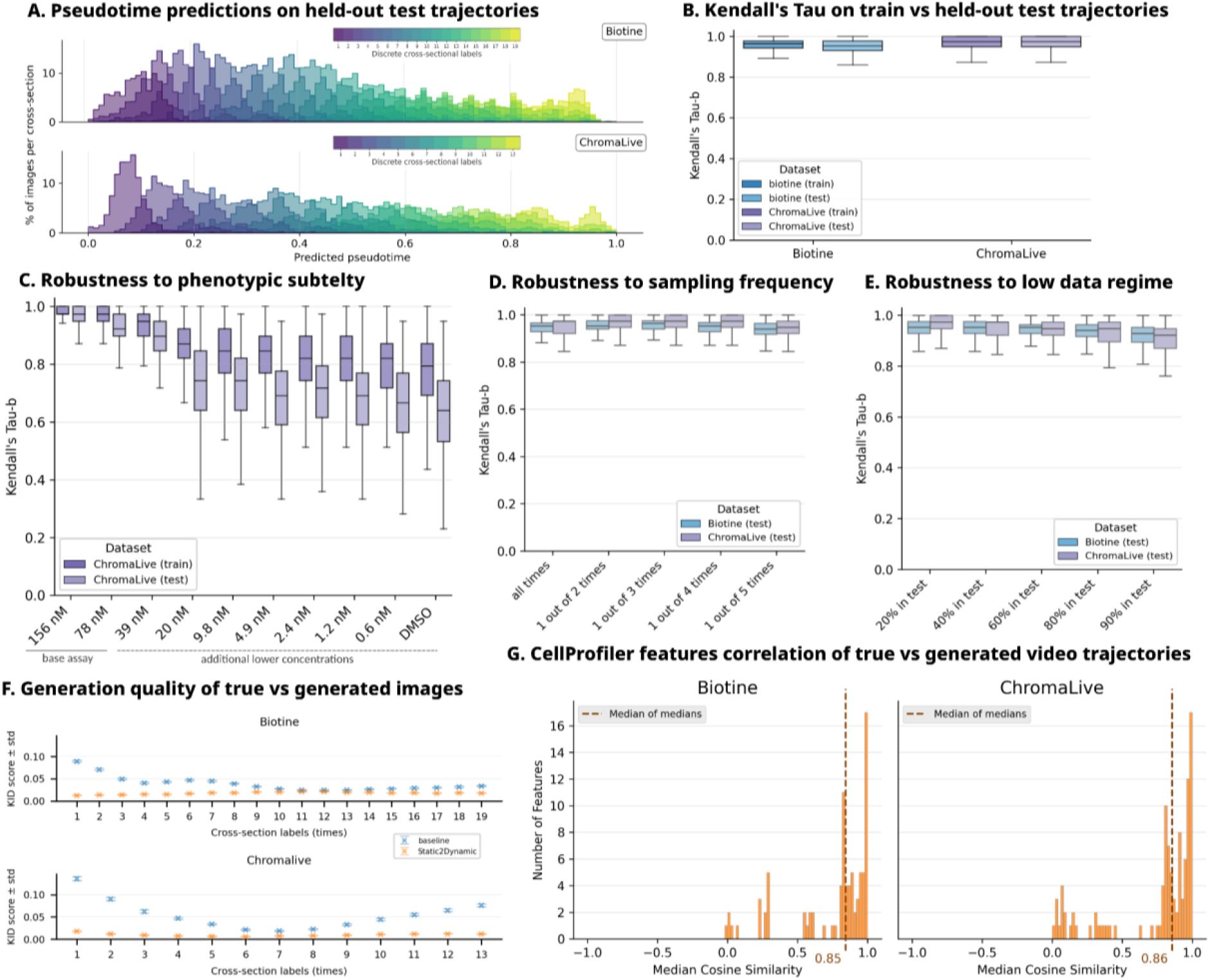
Static2Dynamic evaluations on Biotine and ChromaLive, 2 datasets where the ground truth is available. A - Histograms of pseudotime predictions on the images of the 20% remaining unseen test sequences from the Biotine and ChromaLive datasets, after training the first stage of our method on 80% of these data. B - Kendall’s τ of predicted pseudotime versus real time ordering for Biotine and ChromaLive *sequences*, on train & unseen test sets (0 indicates that the two sequences orders are independent, 1 indicates that the two sequences have the exact same order –see Pseudotime evaluations for details). C - Kendall’s τ of predicted pseudotime versus real time ordering for ChromaLive sequences, for decreasing concentrations of the extended ChromaLive assay (see ChromaLive for details). D - Kendall’s τ of predicted pseudotime versus real time ordering for Biotine and ChromaLive unseen test sequences for decreasing image acquisition frequencies. E - Kendall’s τ of predicted pseudotime versus real time ordering for Biotine and ChromaLive unseen test sequences for decreasing training data size. F - Kernel Inception Distance (KID) between real data and data generated by “Static2Dynamic”. The reference “baseline” is the KID between generated images distribution at time t vs all real images at *other* times. G - Histogram of morphological CellProfiler features trajectories correlations between generated and real videos of the same cell for Biotine & ChromaLive, starting from the same true frame (see Video generation evaluation and **Supp. Fig. 11** for details).

**Figure 3:**
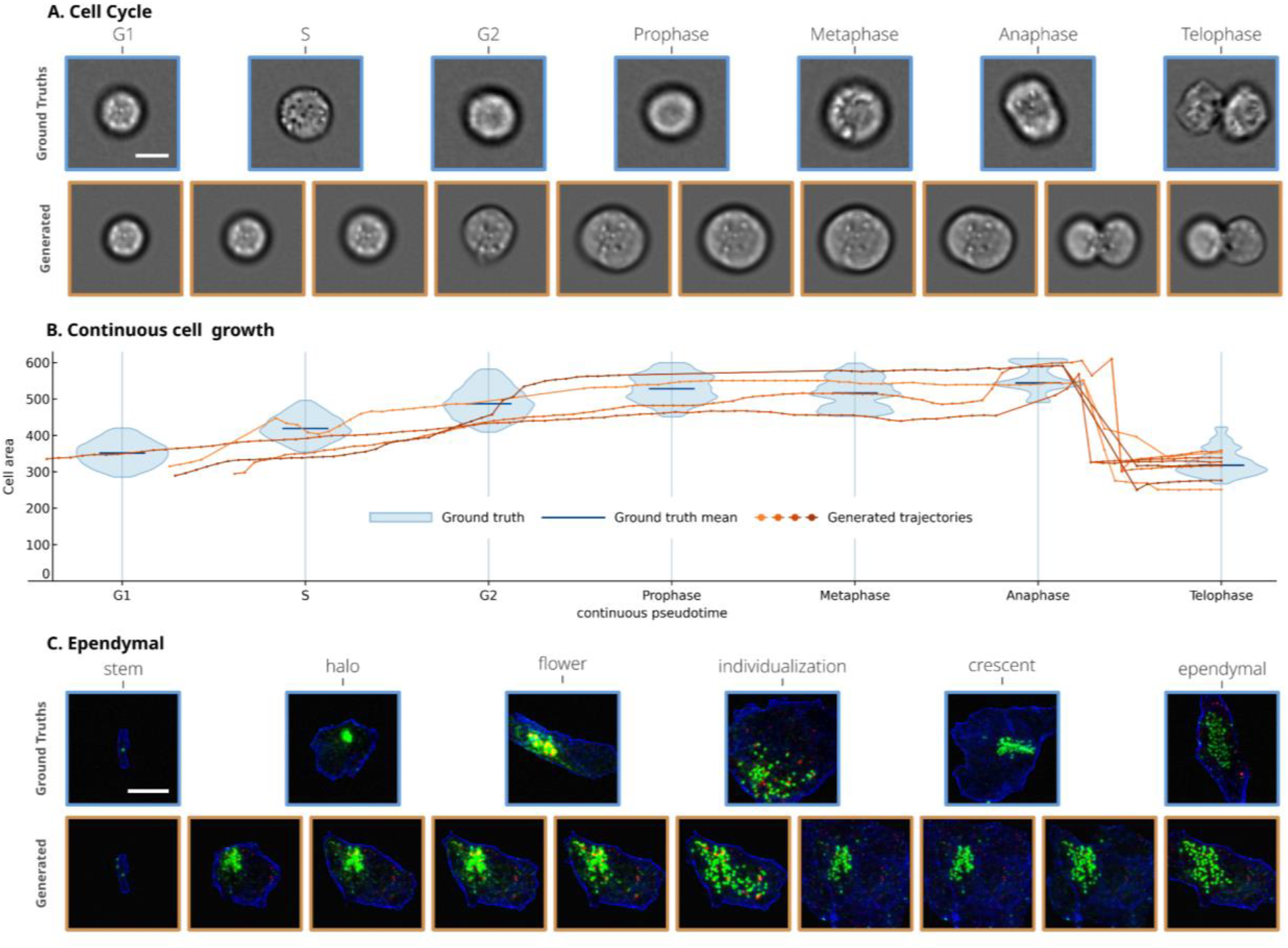
Applications of Static2Dynamic to single-cell processes. A - Reconstruction of single cell cycle video from static imaging cytometry data using cell stages as labels and quantitative analysis over time. In blue, the distribution of cell area is computed on real data for each stage. Static2Dynamic makes possible the continuous computation of cell size (but also of any quantitative feature) on any generated single cell video (orange lines), scale bar is 10 µm. B - Reconstruction of the development of a mouse ependymal cell from ex vivo ependymal wall tissue using differentiation stages as labels. The cell surface delineated by the cell junctions (blue) is gradually growing from a stem cell with a few centrioles (green), centrioles are amplified with the appearance of deuterosomes (red), then the latter disappear. At the end of the process, the ependymal multiciliated cell has many centrioles and gets smaller according to observation, due to its compression by the surrounding ependymal cells in the tissue, scale bar is 20 µm.

Once pseudotime estimation is computed for each image and effectively covers the overall process time range, we leverage it to train a deep image generator model. To this end, we designed a Diffusion Model conditioned on pseudotime embeddings *via* cross-attention (see Methods). The usual denoising objective of diffusion models –specifically, its velocity formulation (Salimans and Ho 2022)– is used to train the neural network on an image reconstruction task, conditional to that pseudotime signal in an end-to-end way. Once completed, this stage provides an image generator that is conditioned on a continuous pseudotime value given as input (**Figure 1**B). Interestingly, the time embedding learnt by the diffusion model showed to be a curve (*aka* a smooth one dimensional manifold) illustrating the continuous nature of the pseudotime representation (**Supp. Fig. 4**).

**Figure 4:**
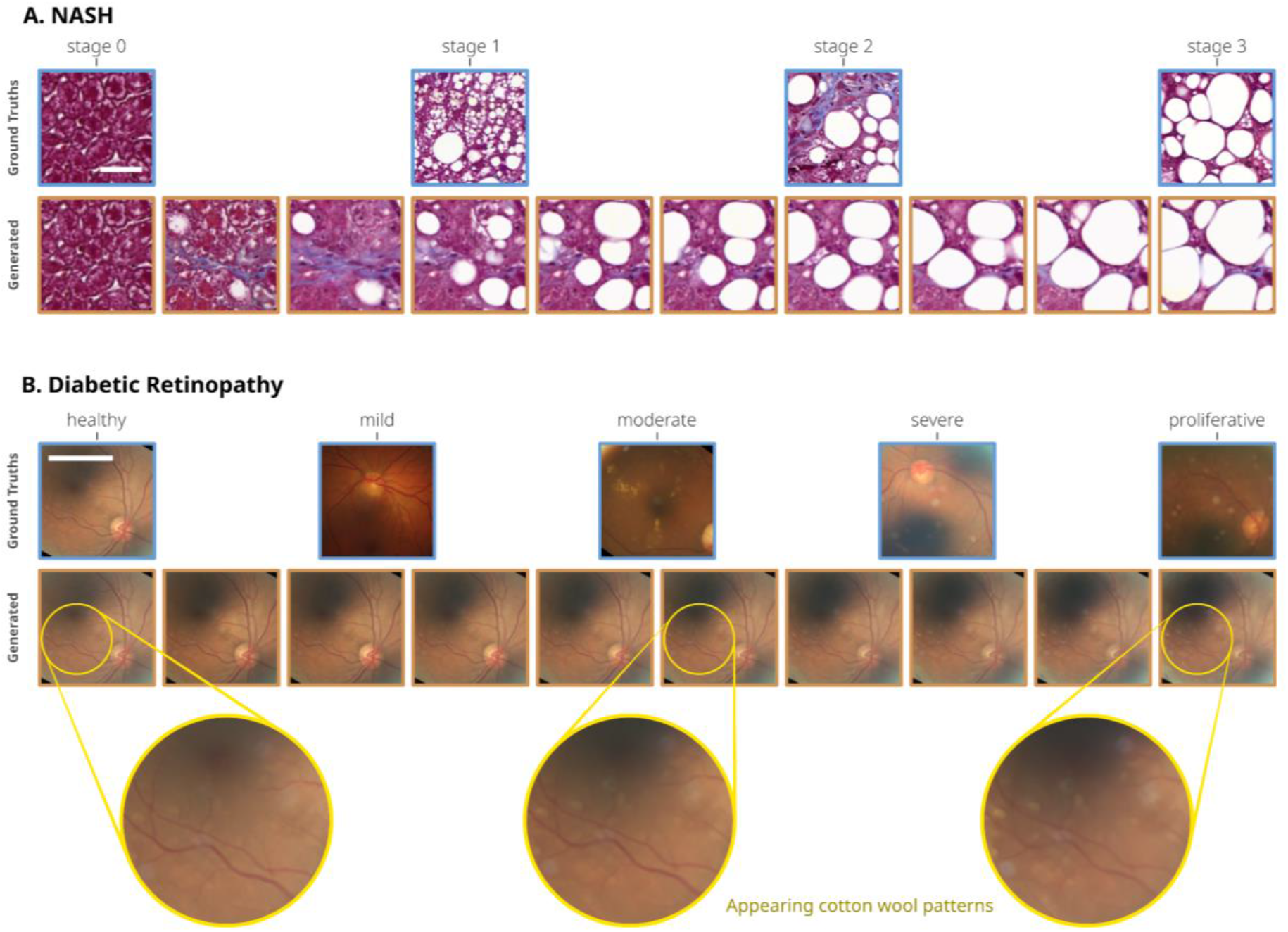
Application of Static2Dynamic to tissue imaging. A - Mouse liver tissue patches at four non-alcoholic steatohepatitis stages (stage 0 is healthy), scale bar is 50 µm. B - Full eye fundus imagery at five stages of diabetic retinopathy. This dataset exhibits highly heterogeneous pixel scales; for reference, the scale bar is about 5 mm.

Generating temporally coherent successive frames of an unseen video sequence of the underlying process can finally be achieved from a single random noise image that is iteratively conditioned with successive pseudotime values from 0 to 1. Interestingly, instead of a random noise image, a deterministic inverted noise can be obtained by DDIM (J. Song et al. 2021) inversion (Dhariwal and Nichol 2021) from any real image of the data distribution, making it possible to animate any real image from the dataset (**Figure 1**C). Static2Dynamic thus offers a way to observe the future of an early stage of the process, or reversely to observe the past of a later stage. Altogether it enables observation of the whole process from any image in the dataset (Videos obtained through this method for all the datasets presented in this work can be found here: biocompibens.github.io/static2dynamic). Interestingly, at first glance our approach removes all statistically unpredictable events from still images such as camera shifts or cell stochastic movements, effectively creating a stable video of the process, free of some of the natural variability found in the empirical data.

### Static2Dynamic accurately and robustly predicts static image pseudotime

To evaluate the robustness of Static2Dynamic, we used two large-scale video datasets. This allows us to compare Static2Dynamic against a known ground truth which is usually not available in real-world applications. The first experiment –”Biotine”– is a time-lapse assay using human A549 lung cancer cells treated with biotin (see Datasets, and **Supp. Fig. 5** or biocompibens.github.io/static2dynamic for examples from the dataset as well as video generations). The second experiment –”ChromaLive”– is a live-cell time-lapse assay monitoring HeLa cells in response to staurosporine treatment (see Datasets, and **Supp. Fig. 6** or biocompibens.github.io/static2dynamic). In both experiments, a single compound treatment effect was monitored over time by live microscopy in many replicates which, once preprocessed, produced respectively 7,680 and 600 (per concentration) 128×128 videos imaging the same global cellular process. For the ChromaLive dataset, the 2 highest concentrations of the compound treatment (out of 10 total) were used as the main dataset and lower concentrations to challenge the robustness of the method. Each training using ChromaLive was therefore performed on a lower number of pictures than with Biotine. In both datasets the cross section labels considered were the acquisition times.

**Figure 5:**
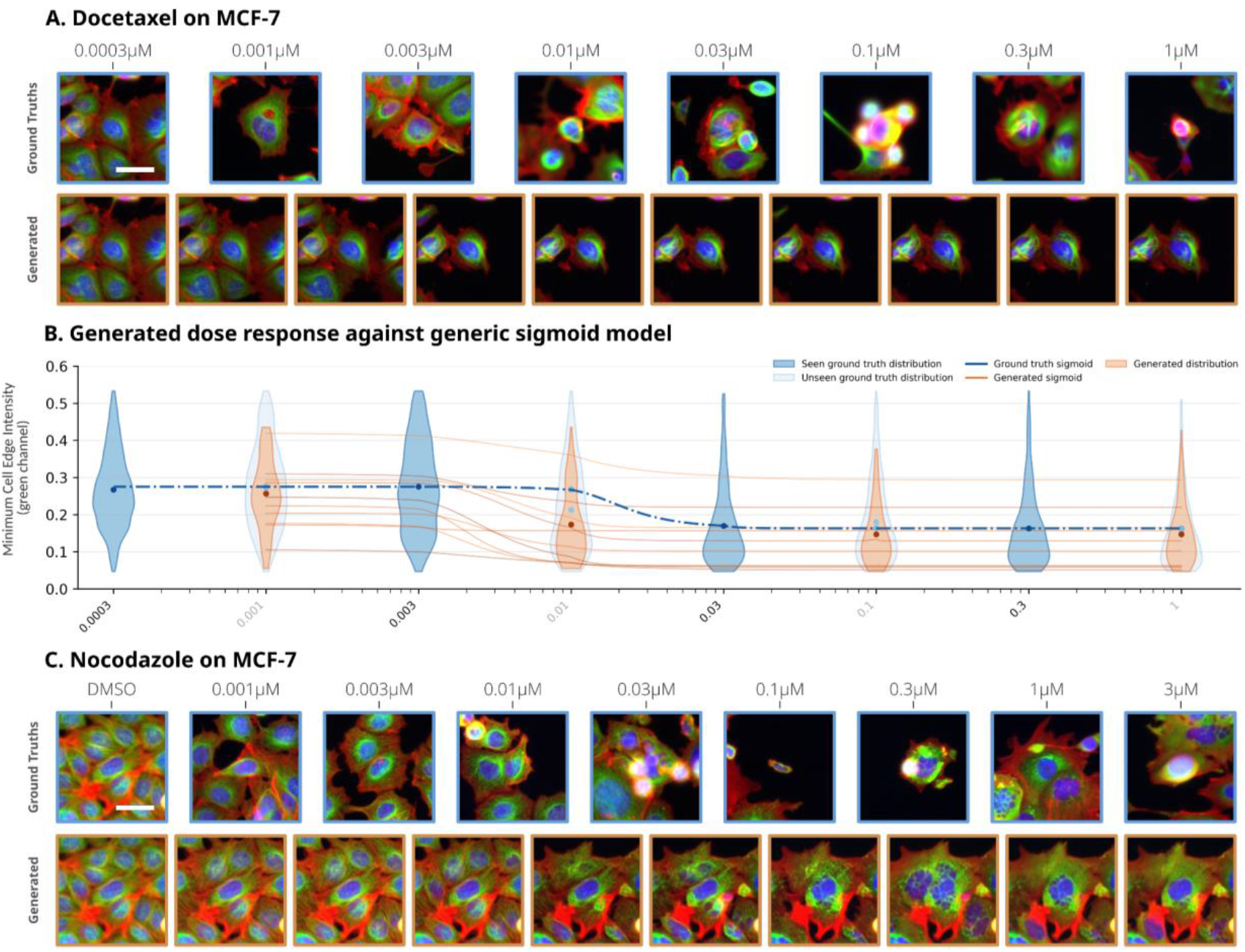
Application of Static2Dynamic to two compounds of a fluorescence drug screening dataset on MCF-7 cells, with increasing concentrations as labels. A - Docetaxel. Static2Dynamic’s generated videos produce phenotypes (orange violins) that closely match the true distributions of Cellprofiler features at unseen concentrations (light blue violins), and the inferred dose responses of generated, *individual trajectories* (orange lines) reveal more precise profiles compared to the *global* response curve inferred from the average real distributions (blue dotted line), notably with a different inflection point than what a sigmoid model on the mean feature values would predict. See **Supp. Fig. 20** for additional ones. B - Nocodazole (with DMSO also included). The scale bars are 30 µm.

To evaluate our pseudotime estimation, which is the first stage of Static2Dynamic, we fitted both the LDA and spline models using images from 80% of the sequences as train set. We then predicted the pseudotimes of the remaining 20% fully unseen sequences of images (**Figure 2**A). We could observe that the estimated pseudotime was consistent with the time arrow. We then compared the real order of images in held out video sequences with the order predicted by our pseudotime estimation pipeline using Kendall’s τ coefficient (Kendall 1938). High Kendall tau scores on both the train and test sets confirmed the precision of our pseudotime prediction (**Figure 2**B). We then assessed the robustness of our method to an increasingly subtle and less visible phenotype by applying it independently on the decreasing concentrations of the ChromaLive assay, down to untreated samples where cells were only affected by time (**Figure 2**C). We further evaluated that this predicted ordering was robust to lower sampling step frequencies (**Figure 2**D) and data scarcity (**Figure 2**E). We also checked that using images from sequences for this evaluation (without providing actual sequence information) did not impair or favor pseudotime estimation (**Supp. Fig. 7**a-d). In addition, we provide evidence through ablation studies that our pseudotime estimation is robust to the choice of the pretrained encoder (**Supp. Fig. 8**). We nevertheless selected DINOv2 for its slightly superior performance for highly variable datasets (**Supp. Fig. 9**). We also provide evidence that the LDA transform step is crucial to emphasize time variant features and recover a reliable trajectory (**Supp. Fig. 10**).

### The generated videos recapitulate unseen real sequence dynamics

To evaluate the accuracy of video generation –the second and third stage of **Static2Dynamic**– at recapitulating dynamic characteristics from the real underlying process, we first evaluated how the continuously conditioned generative model was accurate at reproducing the observed cross sectional image data distributions at each time step. To this end, we estimated the pseudotime of all images in the distribution and generated as many images conditioned by these pseudotime. We then computed Kernel Inception Distances between these image sets for the evaluation datasets Biotine and ChromaLive (**Figure 2**F), but also for all the the other datasets of this paper as it doesn’t necessitate ground truth sequences (**Supp. Fig. 12**. We can observe that Static2Dynamic yields comparable and low KID across all timesteps, ensuring that generation quality is consistently high along the video sequences.

Once generated frame quality had been assessed as correct, we then aimed at performing direct comparison between real video and generated ones for the same cell, in order to evaluate the capability of Static2Dynamic to recover unseen dynamics. However, the generated videos produced by our method are cleared from variations that are not related to the main biological process, such as camera drift, but also stochastic cell displacements that cannot be predicted from static unpaired images only. It can be seen as an advantage because it filters out irrelevant variations that hinder observation of the process of interest, but it stabilizes the view such that pixel to pixel video comparison for evaluation becomes irrelevant. In order to produce a biologically meaningful comparison we thus segmented all cells from both the real and regenerated videos (starting from the same initial true frame), tracked them over time, and finally computed CellProfiler morphological features frame-per-frame, giving us a biologically meaningful signature over time for each true cell feature track and its generated counterpart. By comparing these morphological signatures over time for each true–generated pair of feature tracks, we could evaluate how close a single cell process was recovered at the scale of many whole fields of view (**Supp. Fig. 11**). As such, most CellProfiler features exhibit a strong correlation between the true and generated video tracks (**Figure 2**G).

In the following, we applied Static2Dynamic on multiple and varied biological settings where ground truth sequences are not available.

### Reconstructing the cycle of a single cell from static images

We first applied Static2Dynamic to imaging flow cytometry data that produces snapshot images of single cells. We used manual annotations corresponding to the cell cycle phases as cross section labels (Blasi et al. 2016). Static2Dynamic could reconstruct continuous video dynamics of the cell cycle from these static images, as can be seen in **Figure 3**A (and **Supp. Fig. 13** –also biocompibens.github.io/static2dynamic for a smooth video). Furthermore, we measured that our model successfully recovers cell growth with precision by tracking the size of generated cell videos and comparing it against the static ground truth distributions (**Figure 3**A). Interestingly, thanks to our augmentation strategy, the *continuous* process of division could be recovered from a highly unevenly distributed dataset with very few examples from the Anaphase and Telophase (see **Supp. Fig. 3** and **Supp. Fig. 14**).

### Reconstructing an ependymal single-cell differentiation process

We then applied Static2Dynamic on segmented single-cell from ependymal tissue images acquired by dissecting the brains of developing mice sampled at regularly spaced time (Bankolé et al. 2025). As most developmental processes occurring in utero and lasting weeks, observing in vivo the differentiation of an ependymal cell has remained impossible to date. We first used a manual labelling of a few hundred cells over six different stages of differentiation that were previously identified (Bankolé et al. 2025) as cross section labels to perform our pseudotime estimation on all the remaining images of the dataset, highlighting the capability of our method when a limited number of labels is available (see Ependymal for details). We then performed the last two stages of Static2Dynamic to generate videos of ependymal cell development. We can see, for the first time, the continuous transformation through every stages of differentiation, from stem cell with a few centrioles to fully differentiated ependymal multiciliated cell (**Figure 3**B, **Supp. Fig. 15** and biocompibens.github.io/static2dynamic).

### Reconstructing the nonalcoholic steatohepatitis disease process

We then moved to a larger scale with a dataset of liver tissues of murine rodents suffering from non-alcoholic fatty liver disease (NAFLD)^1^ (Heinemann et al. 2019). Using the four original scores of steatosis annotations provided by veterinary pathologists as cross section labels, Static2Dynamic was able to generate smooth videos of the overall disease process from any image from the dataset (**Figure 4**A, **Supp. Fig. 16** and biocompibens.github.io/static2dynamic). This demonstrates the capability of our method to model disease dynamics at the tissue level despite a low number of initial cross sections.

### Reconstructing the diabetic retinopathy disease process

We then tested Static2Dynamic at the organ level on eye fundus images annotated in five stages of diabetic retinopathy. This challenging dataset (originally a Kaggle classification competition) exhibits subtle phenotypic changes along the disease progression stages and very high imaging variability across samples. Static2Dynamic was nonetheless capable of removing the biological variability in its video generations by keeping the identity of the initial eye fundus, and progressively revealed the characteristic phenotype of the disease so-called cotton-wool spots (**Figure 4**B and **Supp. Fig. 17** or biocompibens.github.io/static2dynamic).

### Using drug concentration as time

Finally, we applied Static2Dynamic to cell painting images of cells treated with 8 concentrations of two compounds: Docetaxel and Nocodazole (Caie et al. 2010). Using concentrations as cross section labels instead of time, we show that not only can Static2Dynamic generate a single cell sequence along a continuous range of intermediate concentrations, but that computing morphological features on such a sequence produces robust individual dose response curves (**Figure 5**B and C, **Supp. Fig. 18, Supp. Fig. 19** and biocompibens.github.io/static2dynamic). Interestingly, single cell dose responses correctly pass through hold out concentrations and avoid the utilization of a parametric model such as a sigmoid fit on average feature values of a cell population. This utilization highlights the general applicability of Static2Dynamic and its ability to model a continuous phenotype intensity variation that is not necessarily related to a temporal process.

## Discussion

We introduced Static2Dynamic, a method designed to reconstruct unseen biological dynamics from cross-sectional image datasets. By leveraging time-unpaired static samples, the approach first infers a continuous pseudotime within a time-discriminative deep representation space, then models the image distribution conditioned on the underlying process, and ultimately generates temporally coherent video trajectories initialized from real observations.

Our extensive evaluation against real video datasets demonstrates that Static2Dynamic can reliably reconstruct biologically meaningful dynamics from purely cross-sectional image data. Across a range of validation settings and concrete applications, our method consistently recovered coherent trajectories and generated plausible intermediate and future states. These findings support the idea that large collections of static images implicitly encode sufficient statistical structure to approximate underlying biological processes, even in the absence of explicit temporal supervision.

A central assumption of the current framework is the existence of a dominant, global process that can be captured as a continuous trajectory in representation space. This assumption holds in many settings and, as a result, stochastic cell displacements and imaging-related variations (e.g., stage drift or focus fluctuations) tend to be filtered out by Static2Dynamic in favor of the dominant trend. This is a clear asset of the method to observe the main process. Furthermore, reintroducing stochasticity into the generative process if needed is straightforward from a technical standpoint, for example by replacing the deterministic DDIM scheduler with a stochastic diffusion scheme. It can provide a more realistic representation of biological variability, particularly in systems where cell fate decisions or microenvironmental fluctuations play a critical role. It could also provide uncertainty estimates of future fates.

While this assumption of a dominant process holds in many settings, it must be acknowledged that it becomes restrictive in more complex biological systems where multiple subprocesses coexist within the same tissue. For instance, events such as cell division, local rearrangements, or rare phenotypic transitions may not correspond to a strong global statistical shift and are therefore effectively ignored by the model. We could identify this limitation on cellular videos presenting mitosis and comparing its reconstruction capability before and after the splitting events (**Supp. Fig. 21**). Cellular systems that exhibit multiple fates and branching behaviors cannot be reduced to a one-dimensional ordering without loss of information. This suggests that explicitly modeling branching dynamics remains a compelling open challenge.

Overall, these limitations point toward a broader question: what aspects of biological dynamics should such models aim to capture? While Static2Dynamic successfully recovers dominant, population-level trends, extending it to account for rare events, branching trajectories and stochastic behaviors will require both richer datasets and new modeling strategies. Addressing these challenges will facilitate moving from descriptive reconstructions toward predictive and mechanistically informative models of biological systems.

Overall Static2Dynamic highlights the possibility of bridging the gap between static snapshots and dynamic biological processes. It enables the inference of both preceding and subsequent visual states from a single image, and allows the simulation of complete trajectories for cellular, developmental, or disease-related phenomena that were not directly observed. It may also find applications where the process is not driven by time but by another continuous variable such as concentration. Such capabilities open new avenues for studying systems where longitudinal imaging is impractical or impossible, opening the door to “pseudotime microscopy”, while also raising important questions about the interpretability, reliability, and biological validity of the reconstructed dynamics.

## Methods

### Probabilistic modelling

Formally, the general problem that we are tackling is well expressed in a probabilistic paradigm: we model the “truth” (ie the distribution of possible videos) as a probability distribution over time-dependent paths in ℝ^*d*^:

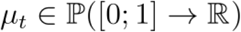

We don’t have access to any realizations of such paths (videos), but rather to collections of static examples at some timesteps: for a given (and problem-specific) finite sequence of cross-section labels *t*_*1<· · ·<*_*t*_*n*_*∈*[0,1], we have 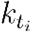observations for each time *t*_*i*_:

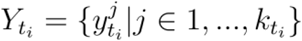

constituting our dataset for that timestep *t*_*i*_.

Our goal is to recover the *whole* dynamic (*μ*)_0 ≦ *t≦* 1_, which in our case is the distribution of videos that could have been acquired from some studied process, had we been able to do so.

The fundamental mismatch between the available data (static images with sparse, discrete labels) and our goal (recovering a coherent video) necessitates continuously distributing training samples along an abstract axis representing process evolution, with the ultimate aim of training a generative model continuously along this axis.

### Pseudotime estimation from unpaired, static image data

The goal of the first stage of our method is to derive a continuous pseudotime label for each sample in the dataset, in order to later condition the image generator on that continuous pseudotime information. To this end, we encode each image in our dataset with DINOv2 (`dinov2-with-registers-giant`) (Oquab et al. 2023), a self-supervised foundational Vision Transformer (ViT) model producing image representations whose interest to Biology has already been well established (Sanchez et al. 2026). The obtained representation, however, does not especially favor variations aligned with the evolution of the process through time (or class labels); note that this first stage is entirely label-agnostic. The goal of this first step is merely to obtain highly discriminative features from our images and we observed that other sufficiently powerful image encoders –such as a generalist convolutional neural network or non-contrastive ViT– could perform reasonably well (see **Supp. Fig. 9** and **Supp. Fig. 8**). To obtain features that are discriminative for the evolution of the process through time, we then leverage the available time labels we have at hand (**Supp. Fig. 1**). These labels might be true timestamps, or gross indication of the stage of evolution of the process. We then projected the DINOv2 embedding onto the subspace obtained with a Linear Discriminant Analysis (LDA) (Fisher 1936; Pedregosa et al. 2012). The LDA has a number of attractive properties: it has a closed-form solution, is cheap to compute, and essentially has no hyperparameters. From **Supp. Fig. 2** It is clear that this transformation produces representations where the axes discriminate time. We then fit an interpolating B-spline (Schoenberg 1946) of order three through the class centroids in the LDA subspace. This spline curve effectively materializes the global one-dimensional evolution of time of the process. In order to then obtain a new continuous pseudotime label for each sample, we project each datapoint onto this spline in the LDA space using the L_2_ distance. We slightly extended the fitted spline beyond the extremal class centroids to account for the most distant points and then rescale the predicted pseudotimes between 0 and 1 (Algorithm 1).

#### Algorithm 1

Static2Dynamic: Pseudotime prediction stage

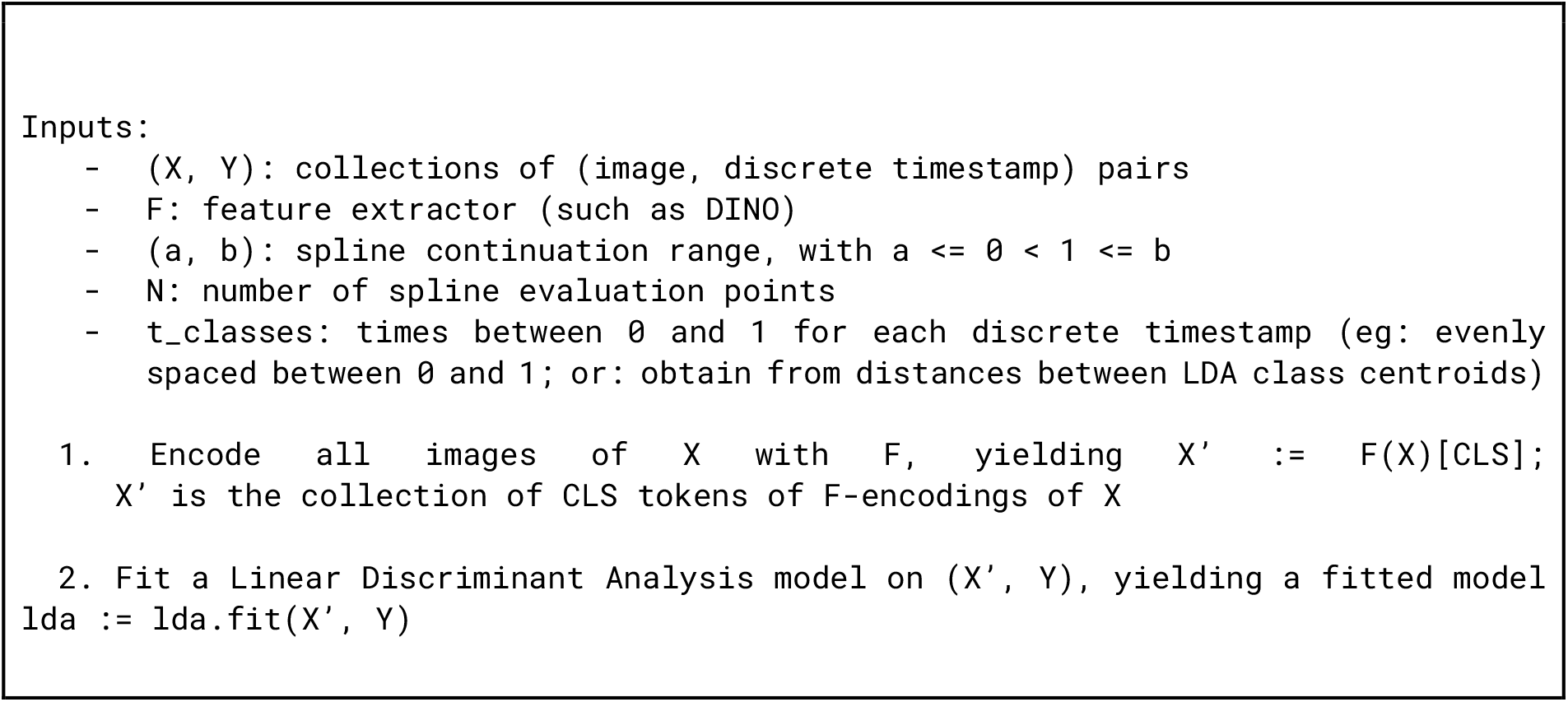
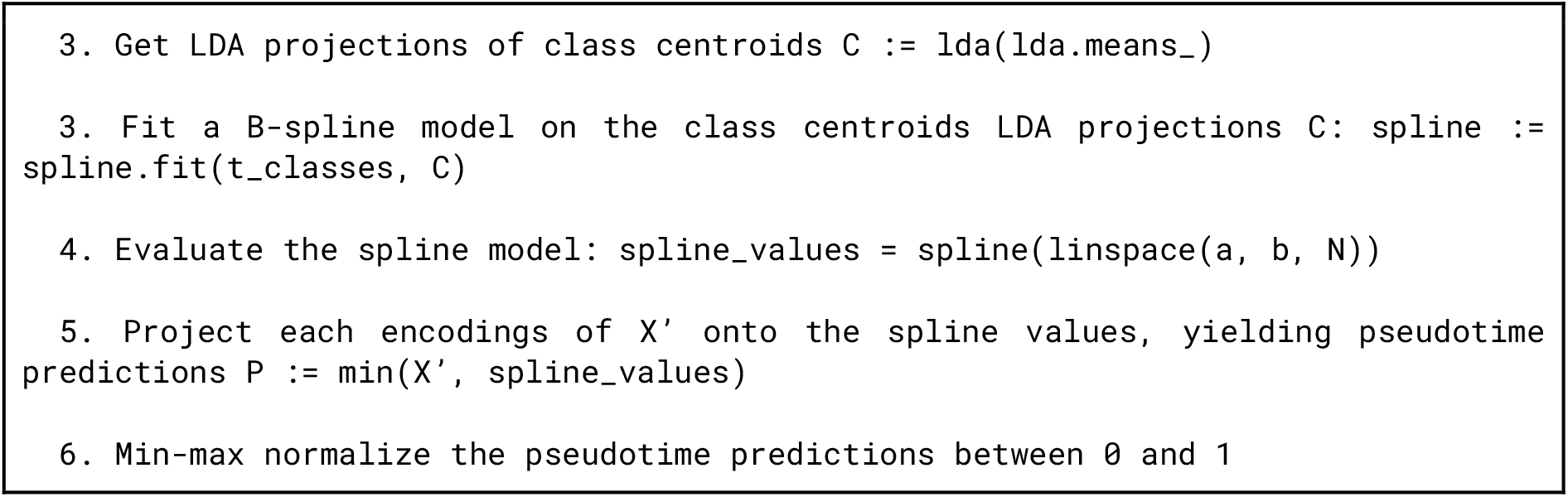

### Diffusion models

Generative modeling consists in creating novel samples from a data distribution to which we only have access through a collection of true examples (the dataset). Recent state-of-the-art methods employed to achieve this task make use of the high expressive power of deep learning architectures and their ability to match very high dimensional data distribution as well as their high degree of controllability. Diffusion Models (DMs) (Sohl-Dickstein et al. 07 2015; Ho et al. 2020; Y. Song et al. 2021), and/or Transformers (Vaswani et al. 2017; Devlin et al. 2018; Radford et al. 2018) are such examples. Generative methods have been applied successfully to many tasks and topics in the scientific domain (Jumper et al. 2021; Abramson et al. 2024; Watson et al. 2023). They have allowed computer simulations to become more reliant on available data, limiting the required explicit modeling prior to some extent. DMs were seminally introduced in a machine learning context by (Sohl-Dickstein et al. 07 2015) and notably popularized in (Nichol and Dhariwal 07 2021) and (Dhariwal and Nichol 2021). They solve the generative problem by modeling the creation of a novel sample as an iterative denoising process, starting from Gaussian noise. More general formulations (Y. Song et al. 2021; Karras et al. 2022) model the noising process as a stochastic differential equation (SDE) of the form:

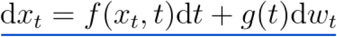

where *x*_*t*_ is a data sample at *t=0* and a sample of the Gaussian distribution when *t* → +∞, and is the Wiener process. Concretely, some final stopping time *T* is chosen to be “large enough”.^2^

There then exists (Anderson 1982) a unique reverse-time process of the form:

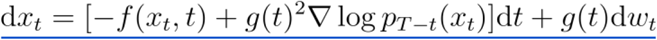

where *P*_*t*_ is the density of the probability followed by solutions of Equation 2. Equation 1 corresponds to denoising data, thus yielding a generative model. Training a DM then consists in learning the score ∇ log*P*_*t*_ over forward trajectories, which is typically done via denoising score matching (Vincent 07 2011).

Deterministic variants of this stochastic formulation have been developed (J. Song et al. 2021) and thorough comparison with not-too dissimilar modellings (Lipman et al. 2022) have revealed the flexibility of the design space of diffusion(-like) models (Albergo et al. 2023; Gao et al. 2024).

### Continuous conditioning on pseudotimes

Allowing the diffusion model to successfully generate samples at different video times necessitates a powerful conditioning scheme. We thus use the same architecture as many successful text/image-to-image/video models and employ a hybrid 2D-ViT/UNet backbone conditioned on video time embeddings *via* cross-attention, where the attention layers are intertwined between ResNet blocks. Specifically, we first embed the continuous pseudotimes obtained from the first stage of Static2Dynamic to sinusoidal encodings, using the usual Transformer (Vaswani et al. 2017) formulation. We then pass these encodings through a small 2-layer MLP projector to learn a relevant time representation (see **Supp. Fig. 4**). Finally, these time representations are forwarded to cross-attention blocks inside the main network (see **Figure 2**B for a schema of our model). The whole system is trained end-to-end using the velocity formulation of the diffusion reconstruction objective (Salimans and Ho 2022). We use AdamW as an optimizer (Loshchilov and Hutter 2019). The stopping decision of the model optimization depends on quantitative evaluations as well as visual checks on generations performed during training. Altogether, after training we obtain an image generator that is smoothly conditioned on pseudotime values from 0 to 1 (Algorithm 2). We leverage this continuous nature of our conditioning for the third and last stage of Static2Dynamic.

#### Algorithm 2

Static2Dynamic: Diffusion Training stage

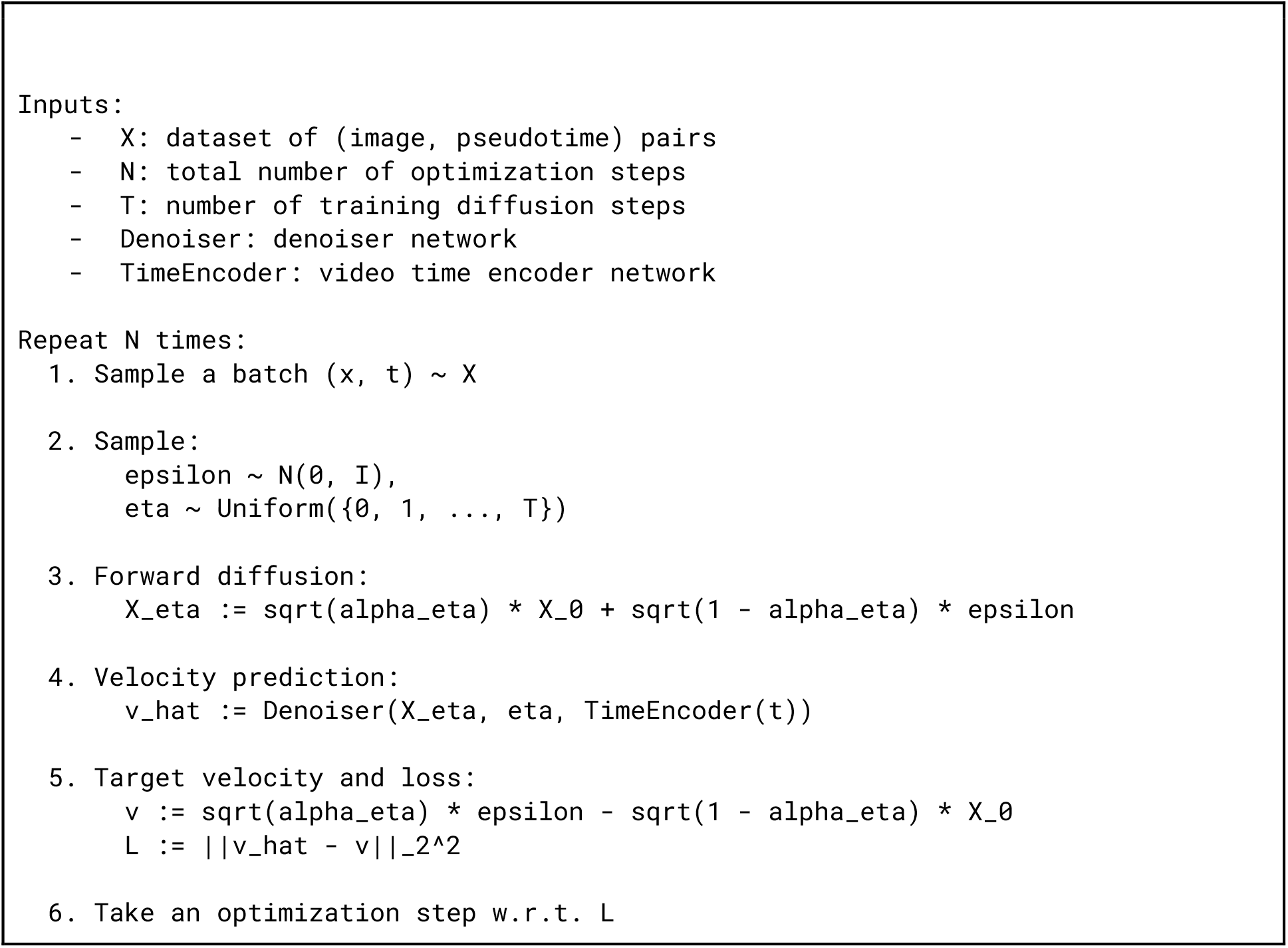

### Video generation

From the previous stage we obtain a model capable of generating images conditioned to a pseudotime. In order to root the generations of the model to a real image of which we want to obtain a future or past evolution, we leverage the DDIM formulation of diffusion models (J. Song et al. 2021). DDIM allows for 1) accelerated sampling and 2) simple inversion of a starting image into the Gaussian sample that would be regenerated back into that very image. This technique, called inversion, is essential for our system as we use the inverted Gaussian sample as the latent representation from which we generate a whole video. An interesting property arises from our usage of the DDIM setting: as the transport learned by our model under this formulation appears to be close to the (regularized) optimal one (Su et al. 2023) (although not equal to it: (Lavenant and Santambrogio 2022)), it indeed produces “optimal” generation in the sense that it tends to remove any stochastic behavior in the video, thus removing the undesirable natural variability and instead effectively creating a pure phenotypic variation over time as video (Algorithm 3).

#### Algorithm 3

Static2Dynamic: Inference stage

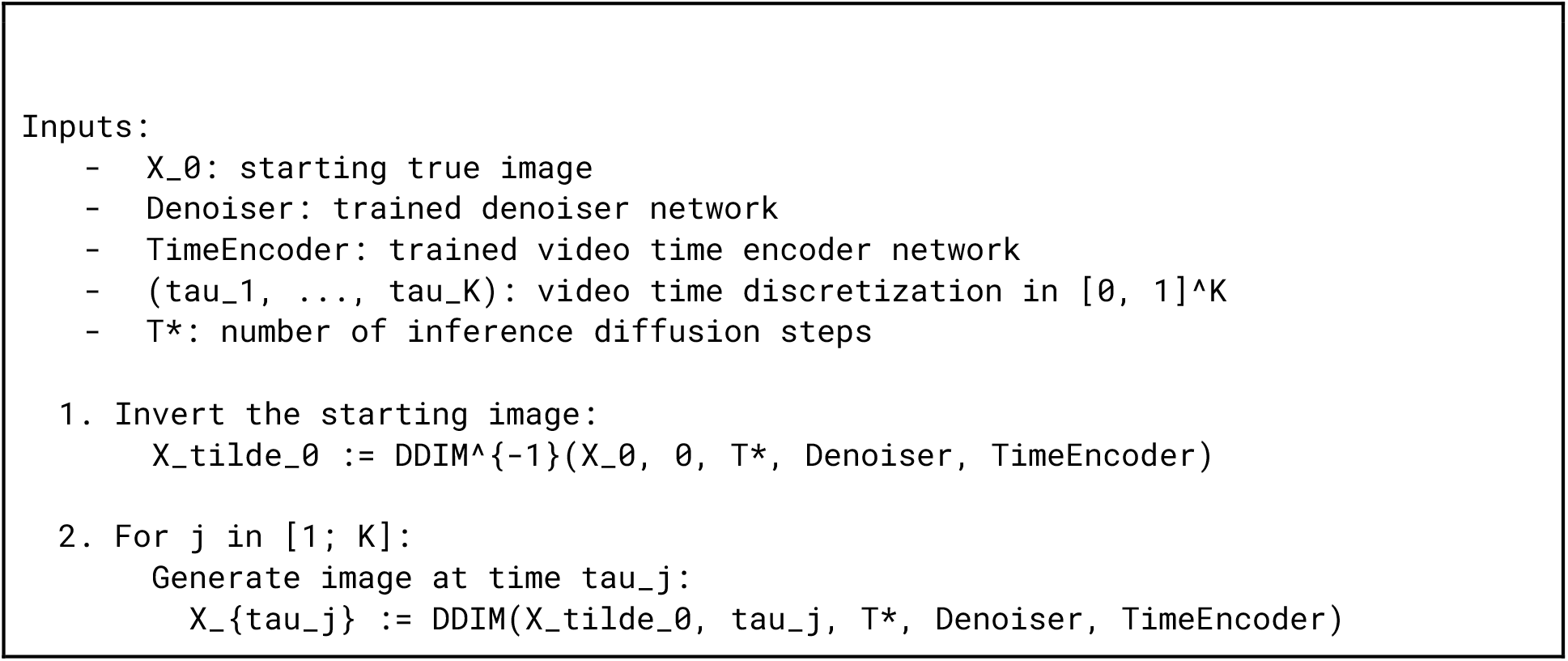

### Pseudotime evaluations

In order to evaluate the fundamental capability of Static2Dynamic to recover the unseen dynamics from static, unpaired snapshots of data, we performed tests at multiple levels: on the predicted pseudotime obtained from the first stage of the method, on the generate images after training the second stage, and on the generated videos from the third and last stage. To perform these evaluations we use our 2 video datasets (biotine and ChromaLive), on which we can compare our generations to actual ground truths.

We started our evaluations by assessing whether the pseudotime obtained from the first stage of Static2Dynamic was correctly predicted. Indeed, the original *ordering* of time labels of the two video datasets can be regarded as perfect information ; we thus expect the model to avoid any reordering of the predicted pseudotime vis-à-vis that original time labels ordering of individual sequences.^3^ To measure this, we employ Kendall’s tau statistics (Kendall 1938, 1945). It is generally expressed as:

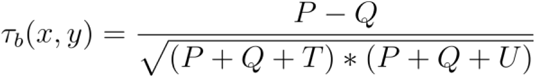

where:

-*x* and *y* are 2 sequences of scalar values

-*P* is the number of concordant pairs

-*Q* the number of discordant pairs

*-T* the number of ties only in *x*

*-U* the number of ties only in *y*

Note that in our case it reduces to:

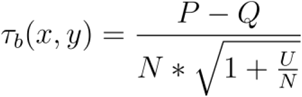

where:

-*x* is a sequence of ground truth labels

-*y* is a sequence of pseudotime predictions

-*P* is the number of concordant pairs

-*Q* the number of discordant pairs

-*N* = *P* + *Q* is the total number of pairs since there are no ties in ground truth time labels. This correlation coefficient approaches 1 as the ranking of predicted pseudotimes is well aligned with that of the original time labels.

We evaluated the pseudotime predictions of Static2Dynamic on different settings (**Figure 2**A-E), always using real trajectories of fully held-out fields as test data. In **Figure 2**B we checked whether the pseudotime predictions on these held-out test trajectories (see **Figure 2**A for their histogram) exhibit good Kendall Tau scores compared to the tau statistics obtained on training data.

Next, in **Figure 2**C, we tested whether Static2Dynamic is capable of correctly ordering samples with decreasing concentrations, down to pure DMSO medium. To this end, we trained the first stage of our method separately on each concentration of the ChromaLive assay (yielding a phenotype that gradually becomes more subtle with the decreasing concentration, **Supp. Fig. 22**), and reported Kendall tau score on held-out data of the same concentration. In **Figure 2**D and E we evaluated the robustness of the first stage of Static2Dynamic w.r.t. a low number of time labels or a low data regime. These evaluations are important because they correspond to real applications (e.g. the NASH dataset has only 4 stages of annotated disease progression, while the Ependymal and Diabetic Retinopathy datasets are significantly unbalanced between classes). For this purpose, in **Figure 2**D, we removed time labels and their associated data at an increasing frequency for both evaluation datasets, trained the first stage of our method on each of these sparse-time dataset variants, and reported the Kendall-tau score for unseen test trajectories. We performed the same kind of stress-test in **Figure 2**E but instead of removing entire times steps we increased the fraction of held-out fields of view in the test split (thus decreasing the amount of data available for training).

### Video generation evaluation

We first report in **Figure 2**F Kernel Inception Distances (KIDs) (Bińkowski et al. 2018) of true vs generated images, for each timestep of the biotine and ChromaLive datasets. KID is akin to FID (Heusel et al. 2017) but less sensitive to small sample sizes, an important feature in our setting. It measures the dissimilarity between the true and generated image distributions by computing the squared Maximum Mean Discrepancy of InceptionV3 (Szegedy et al. 2016) embeddings of both images distributions. For each class, the generated dataset was obtained by conditioning the generation on the distribution of the predicted pseudotimes of the true images of that class (**Supp. Fig. 12**).

We then evaluated Static2Dynamic on its ability to recover the *unseen dynamics* of ground truth data. To this end we generated videos starting from ground truth initial frames of both Biotine and ChromaLive datasets, segmented each cell along time, and computed CellProfiler (CP) (Stirling et al. 2021) features of these single-cell video cutouts. We then compared the obtained CP time-indexed feature vectors of each true versus generated cell pair by computing the cosine similarity between their feature vector, for each feature. After removing highly correlated features, we also removed features that did not show significant variations along time to avoid including high cosine similarities of static features (see **Supp. Fig. 23**).

We then report the histogram of median cosine similarities across all cell pairs, for each feature in **Figure 2**G. There are 6381 true-generated pairs of trajectories across 15 fields for biotine, and 563 pairs of trajectories across 24 fields for ChromaLive. We also report the median of all these medians (across all features) in red vertical dashed lines. Details of the full pipeline can be found in **Supp. Fig. 11**.

We also performed an analysis on mitosis events that happened in the true videos. To this end we grouped pairs of trajectories based on whether a split was detected in the true trajectory (see **Supp. Fig. 21**).

### Data augmentation

A typical challenge in machine learning is data scarcity. Biological imaging datasets often fall into the low data regime due to the complex and expensive nature of the data acquisition. Static2Dynamic is particularly prone to this challenge as its first stage ends up *spreading* samples along a continuous pseudotime axis. We have observed that low data density regions along that axis naturally cause the model to collapse and memorize the sometimes very few available samples. To mitigate this effect, we heavily rely on classical data augmentation techniques, leveraging the invariances of the data. As such, by default, all datasets make use of on-the-fly 8x augmentations of the degree-4 dihedral group. For the Diabetic Retinopathy, Cell Cycle, and Ependymal datasets, we instead make use of more aggressive augmentations by rotating images continuously around the circle. Finally, for the ChromaLive dataset, we train Static2Dynamic on randomly sampled crops over the full fields of view.

### Datasets

#### Biotine

A time-lapse assay was performed using human A549 lung cancer cells (ATCC, CCL-185) stably expressing a GFP-tagged RUSH (Boncompain and Perez 2013) reporter and a hook protein. This system allows for the biotin-induced synchronous release of secretory cargos from the endoplasmic reticulum. Cells were seeded in 384-well plates (Viewplate 384, Perkin Elmer) and maintained for 24 hours prior to treatment. Cells were treated with 40 µM of biotin to initiate cargo trafficking. Time-lapse imaging was performed using the INCell 6500HS automated imaging system (Molecular Devices) at 20× magnification (Nikon Plan Apo, 0.45 NA). To capture the dynamic redistribution of the GFP-cargo, images were acquired across three channels including NucLight for nuclei and rhodamine phospholipids for cell boundaries over a total duration of 90 minutes.

A total of 7,680 255×255 videos were then cropped from the regular grid of the 120 fields-of-view movies. These videos were then rescaled to 128×128.

While we have previously described this system for static high-content screening transfer learning analysis (Corbe et al. 2023), we adapted the protocol for continuous live-cell imaging to provide a dynamic ground truth for Static2Dynamic.

#### ChromaLive

A time lapse assay from (Lippincott et al. 2025) HeLa cells were exposed to a panel of ten staurosporine concentrations ranging from 0 to 156.25 nM. Staurosporine, a broad-spectrum protein kinase inhibitor, is known to induce apoptosis through activation of caspase-3. Following treatment, cells were subjected to the Live Cell Painting assay (ChromaLIVE™). Time-lapse imaging was then performed using spinning-disk confocal microscopy on a Yokogawa CQ1 system, with images acquired every 30 minutes across three z-planes per field of view over a total duration of six hours.

Apart from the study in panel C of **Figure 2**, only the 2 highest concentrations were used (156 and 78 nM). These 2 concentrations correspond to 24 fields-of-view movies, from which 48,000 380x380 videos were cropped *at random coordinates and orientations* (2,000 crops per fields-of-view movie). These videos were then rescaled to 128×128.

For the study in panel C of **Figure 2**, we instead used all 120 fields-of-view movies, from which 48,000 380×380 videos were cropped *at random coordinates and orientations* (400 crops per fields-of-view movie). These videos were then rescaled to 128×128.

#### Cell cycle

An imaging flow cytometry dataset of Jurkat cells from (Blasi et al. 2016). The discrete classes are annotated cell cycle phases (G1, S, G2, Prophase, Metaphase, Anaphase, Telophase). Although a version of this dataset can be found on Broad Bioimage Benchmark Collection (Ljosa et al. 2012), we reprocess the raw data kindly provided by the authors. We only use the brightfield channel and standardize the single cell crops to squares, fulfill the background and align the intensity. A total of 32,266 66x66 images are available in this dataset.

#### Nocodazole and Docetaxel on MCF-7

These images are subsets of the BBBC021 dataset (Caie et al. 2010), available from the Broad Bioimage Benchmark Collection (Ljosa et al. 2012). It is a phenotypic profiling assay of MCF-7 breast cancer cells treated with a collection of compounds over 8 concentrations each. The cells are labeled for DNA, F-actin, and B-tubulin. We use 2 compounds (independently): Nocodazole and Docetaxel. A total of 15,843 and 4,793 196×196 images are available in these datasets, respectively

#### Ependymal

These images were obtained from a previous study on ependymal cells (Bankolé et al. 2025). Lateral ependymal wall tissues were collected at multiple postnatal stages of mice development and processed for immunofluorescence staining. High-definition images of ventricular whole-mounts comprising labeling of cell junction (β-catenin), centrioles (FOP or centrin), and deuterosomes (deup1 or sas6) were acquired. Imaging was performed across all developmental stages to capture the spatial and temporal distribution of these markers. 2D images of the ventricular surface were extracted from the 3D stack (Shihavuddin et al. 2017). Cells were segmented and pasted in empty individual images to build the dataset. A total of 26,310 256x256 images are available in this dataset, of which 4,888 are labelled.

#### NASH steatosis

We use a preprocessed dataset from (Heinemann et al. 2019) consisting of tissue sections of mouse or rat liver tissues suffering from non-alcoholic fatty liver disease (NAFLD, now known as Metabolic dysfunction–associated steatotic liver disease) and stained with Masson’s trichrome. The data is then annotated for 4 stages of steatosis (excess of fat) corresponding to the so-called Kleiner score (Kleiner et al. 2005). A total of 4,571 images are available in this dataset.

#### Retinopathy

We repurposed a challenging Kaggle classification dataset (“Diabetic Retinopathy Detection” 2015) of high-resolution retina images (or eye fundus) with a high variance in the imaging conditions, and subtle phenotypic differences. The data has been annotated for 5 scales of Diabetic Retinopathy (DR): No DR, Mild, Moderate, Severe, Proliferative DR. A total of 35,126 images are available in this dataset.

## Data availability

The following datasets are publicly available via their original research publications: ChromaLive, Cell cycle, NASH steatosis, Retinopathy, Nocodazole and Docetaxel on MCF-7 (see the Datasets section). Biotine and Ependymal are in-house datasets. They are available upon reasonable request.

## Code availability

The code of Static2Dynamic can be found at github.com/biocompibens/static2dynamic.

## Acknowledgements

This work has received support under the program *Investissements d’Avenir* launched by the French government and implemented by the ANR, with the references: ANR-10-LABX-54 MEMO LIFE ANR-11-IDEX0001-02 PSL* Research University, TB was funded by Inserm ITMO Cancer 2023 - DYNAMO. This project was provided with AI computing and storage resources by GENCI at IDRIS thanks to the grant 2024-AD011014994 on the Jean Zay supercomputer’s H100 partition. We thank Elodie Anthony (Institut Curie/BioPhenics) for technical assistance, Franck Perez (Institut Curie/UMR144) for providing the RUSH cell line model, Ibrahim Bilem from Sagaro Bioscience for the ChromaLive dataset, Mathieu Blasi for providing us with the raw data of the Cell Cycle dataset, the computing service of IBENS and Mary-Ann Letellier for proofreading the manuscript.

## Author contribution

T.B. and A.G. proposed the method. T.B. implemented the method and performed all processing and analysis. E.D.N. designed and imaged the Biotine assay, N. S. designed and imaged the Ependymal dataset. A.G. and TB wrote the manuscript. All authors edited the manuscript.

## Footnotes

1 Now known as Metabolic dysfunction–Associated steatotic liver disease (MASLD).

2 Notice that whether T is “large enough” or not depends on the actual noise schedule g that is chosen. Recent works (Albergo et al. 2023) have logically alleviated this artificial pseudo infinite time.

3 In the case of complex processes such as disease progression, we might generally expect some level of discrepancy between samples regarding their ground truth labelling, but time labels of video dataset are perfect ground truth. And even though there might still be discrepancies in the effect of the compound between different well replicates for example, we nonetheless expect that, given a field of view, the state of the process that we want to model (the compound effect) can only be *increasing* with time. So the *ordering* of predicted pseudotimes should never be changed w.r.t. the original ordering of time labels (ie: always increasing).

