## Supplementary Information for "Static2Dynamic: Reconstructing videos of unobservable cellular, developmental, and disease processes"

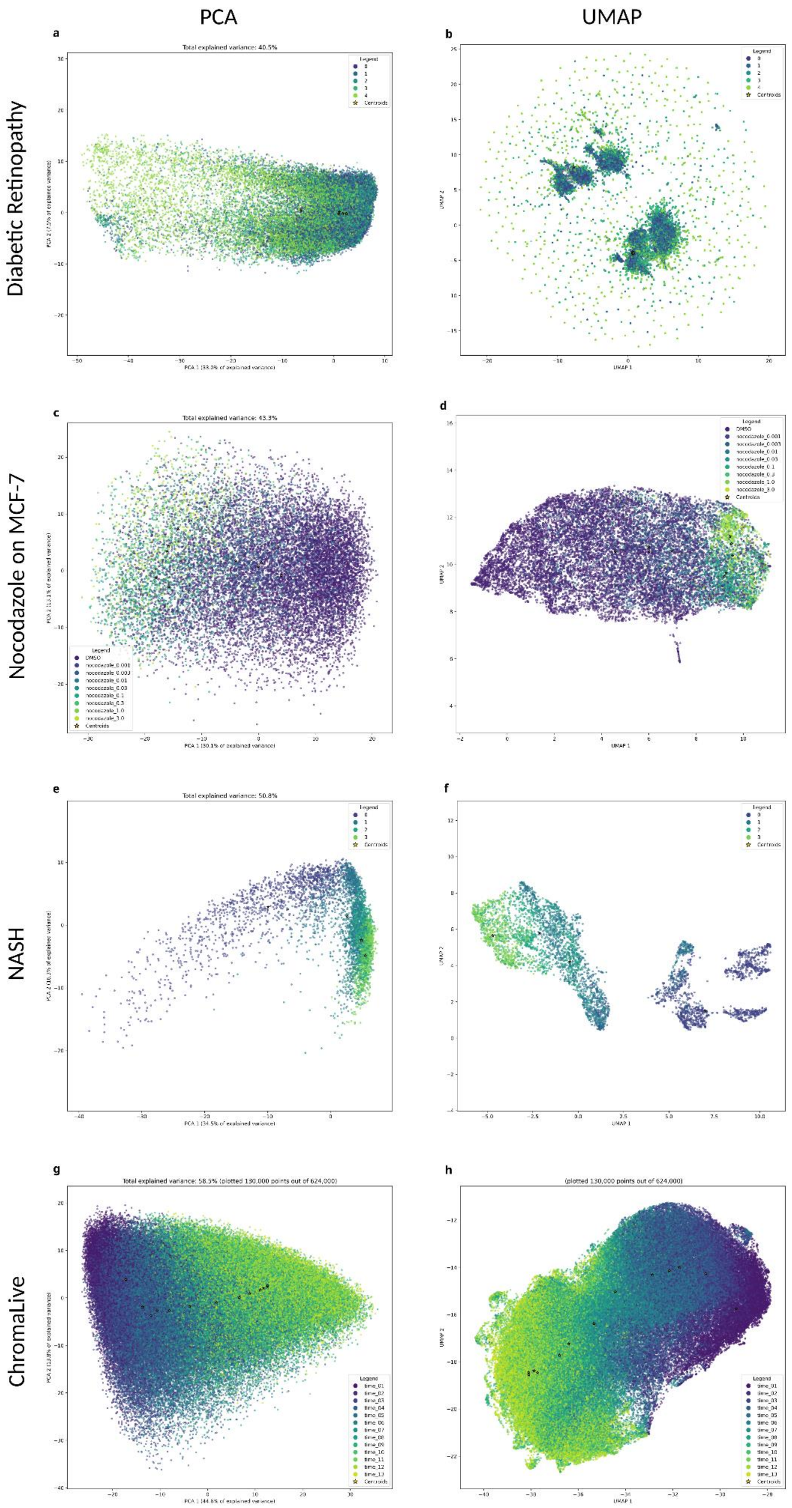


**Supplementary Figure1: PCAs or UMAPs do not directly emphasize time variations for most datasets** PCA (left column) and UMAP (right column) of 4 datasets: Diabetic Retinopathy (top row), Nocodazole BBBC021 (second row), NASH (third row), and ChromaLive (bottom row). Colors are the cross section labels for each dataset.

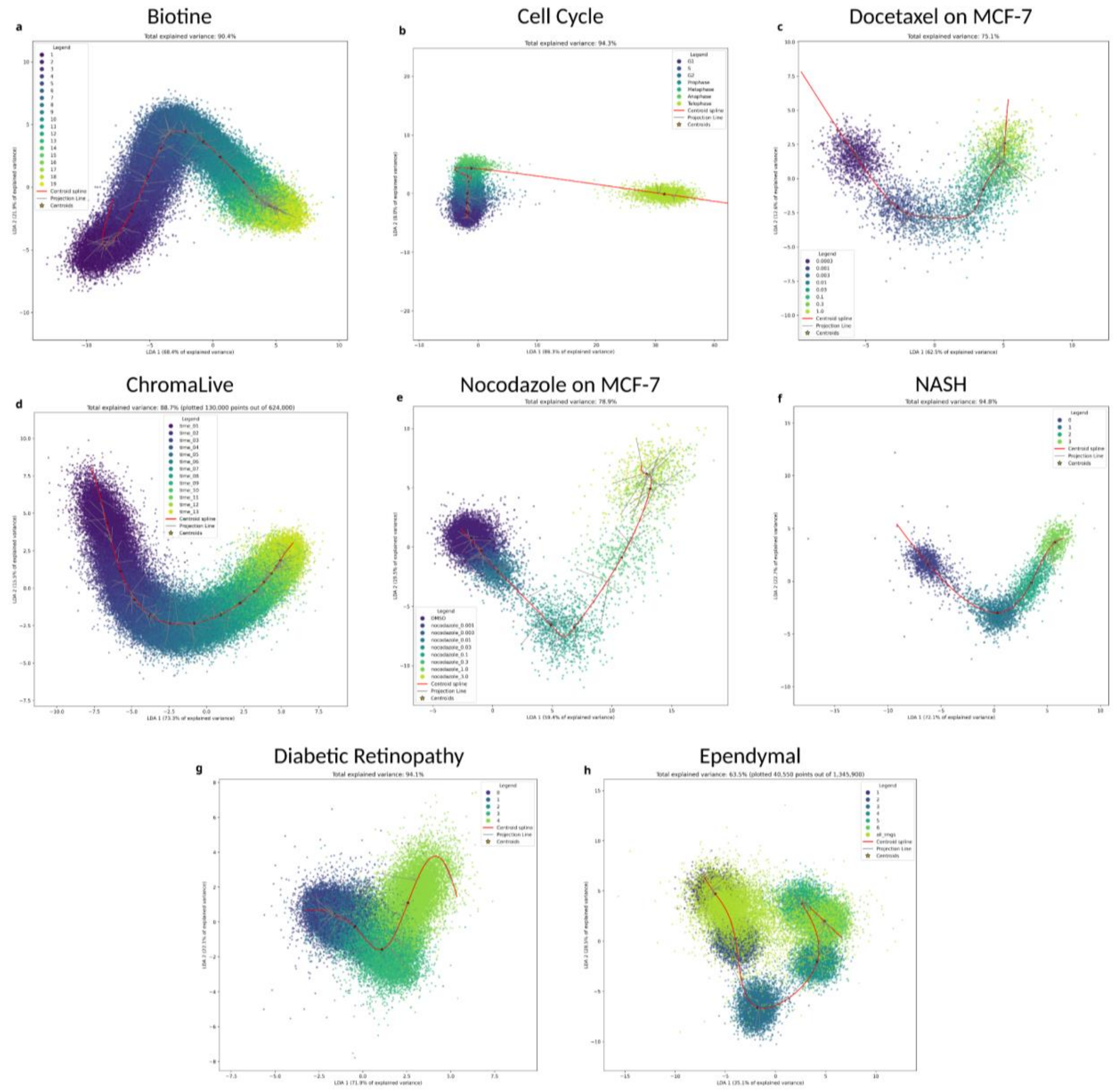


**Supplementary Figure2: Fitted splines in LDA subspace enable us to see and extract a clear general time evolution trajectory on all datasets**. Shown here are the first 2 dimensions of the LDA projections, for all datasets, with fitted B-splines shown in red and some projections from samples to the curve in gray.

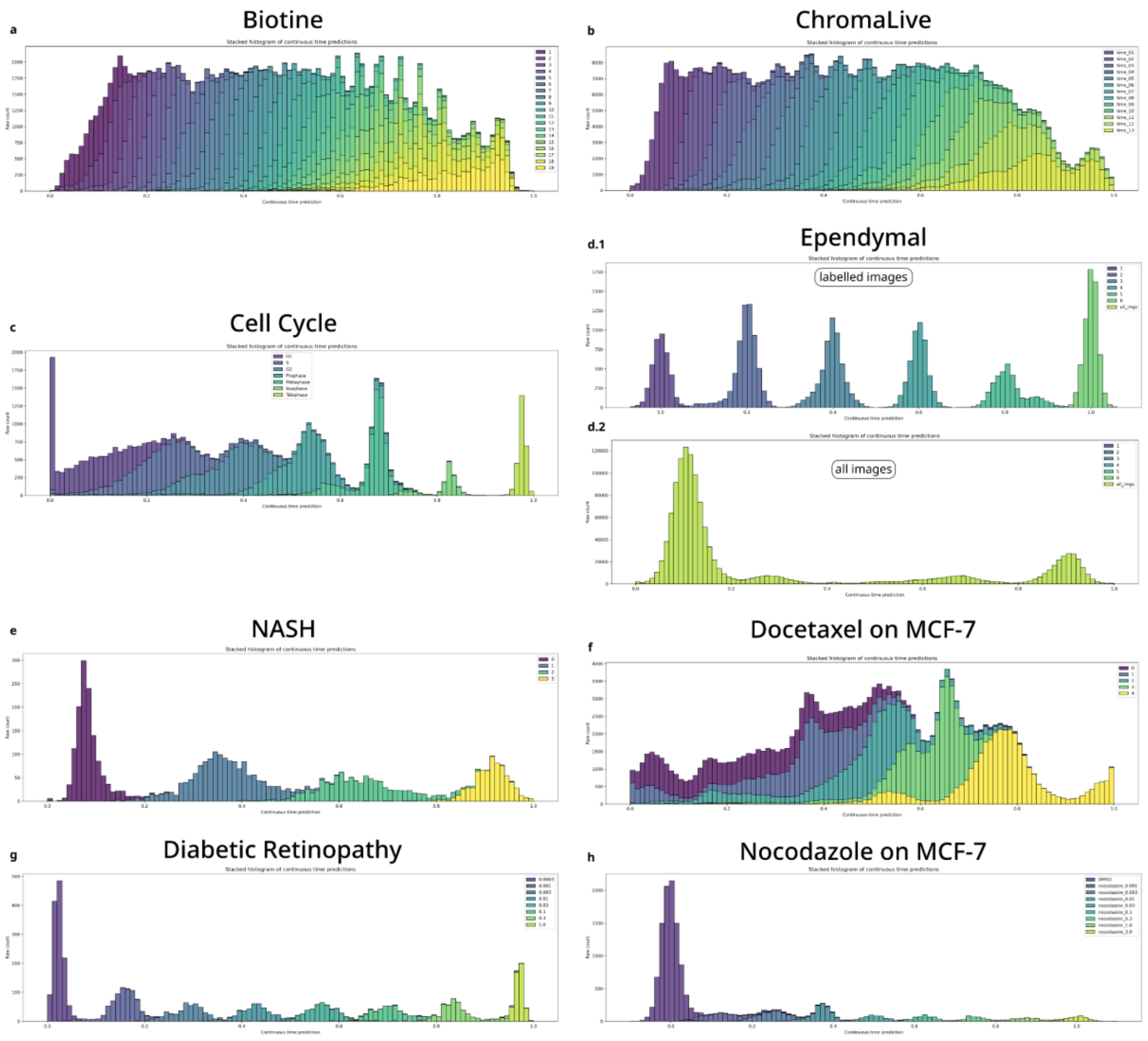


**Supplementary Figure3: Global consistency of predicted pseudotimes.** Shown here are stacked histograms of the predicted pseudotimes for each cross section label, for all datasets. Static2Dynamic preserves the global time arrow given by these cross sections while reordering individual labels when applicable. Two histograms are shown for the Ependymal dataset: one for manually annotated images (4,888 in total), and one for all the remaining images that were not annotated (21,422 images). Note that image counts shown here include data augmentation.

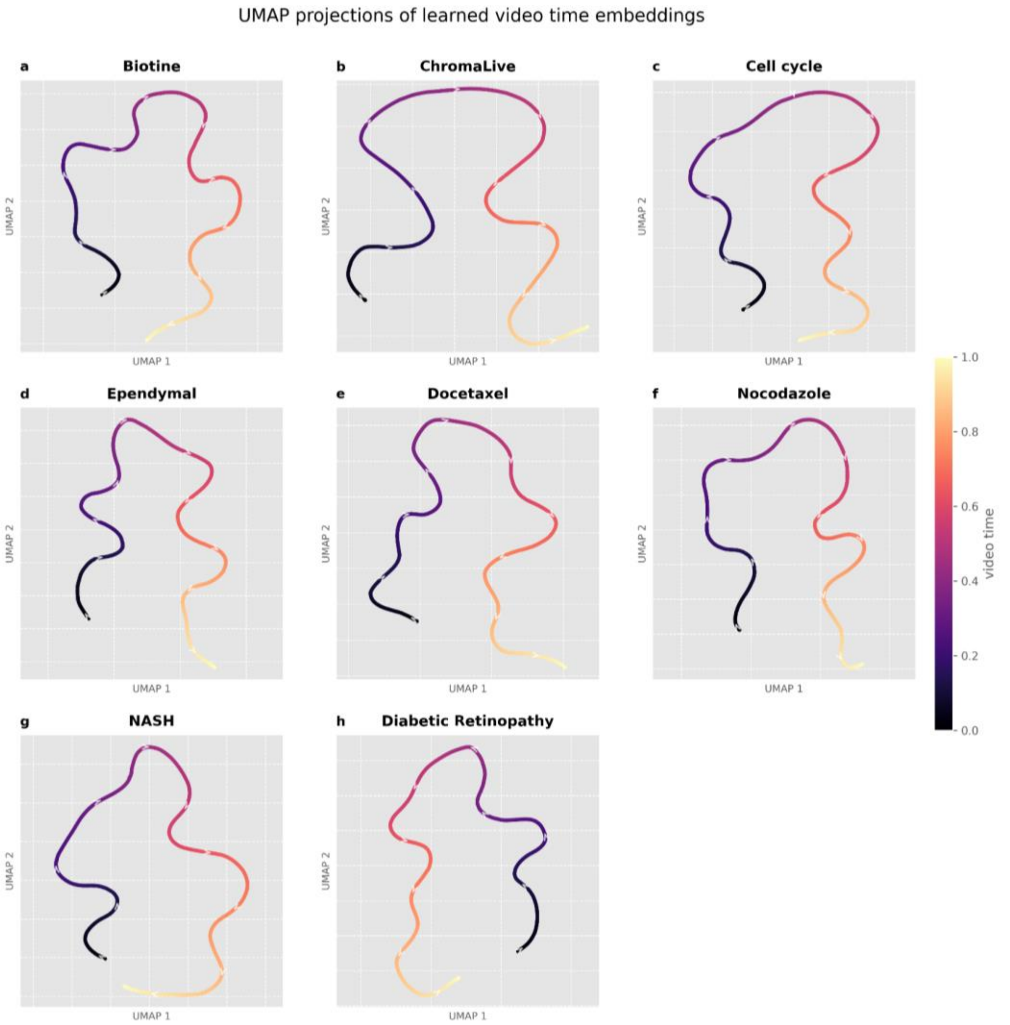


**Supplementary Figure4: Continuous nature of the pseudotime representation.** The time embedding learned by the conditional diffusion model of Static2Dynamic appears as a smooth curve on all datasets.

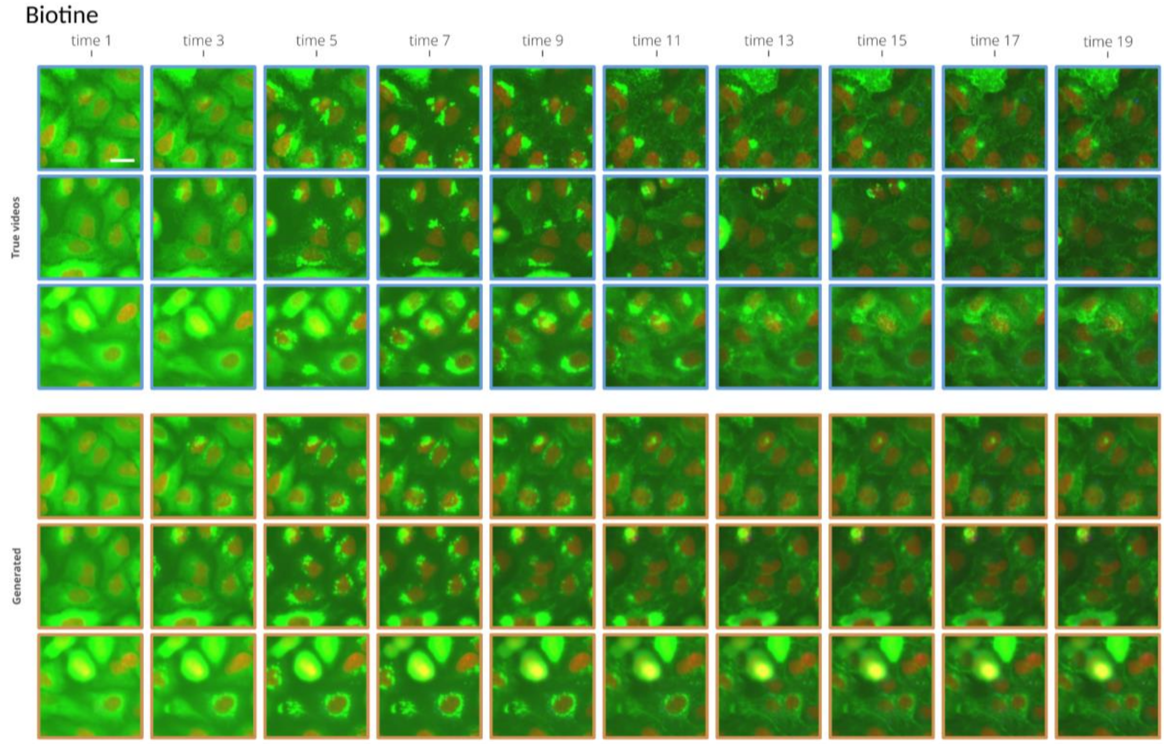


**Supplementary Figure5: True and generated videos for Biotine.** Columns: subsampled video time steps. Top 3 rows: 3 true videos. Bottom 3 rows: 3 generated videos from the same first image. While the true video shifts toward the bottom, the generated video remains stable. The scale bar is 20 µm.

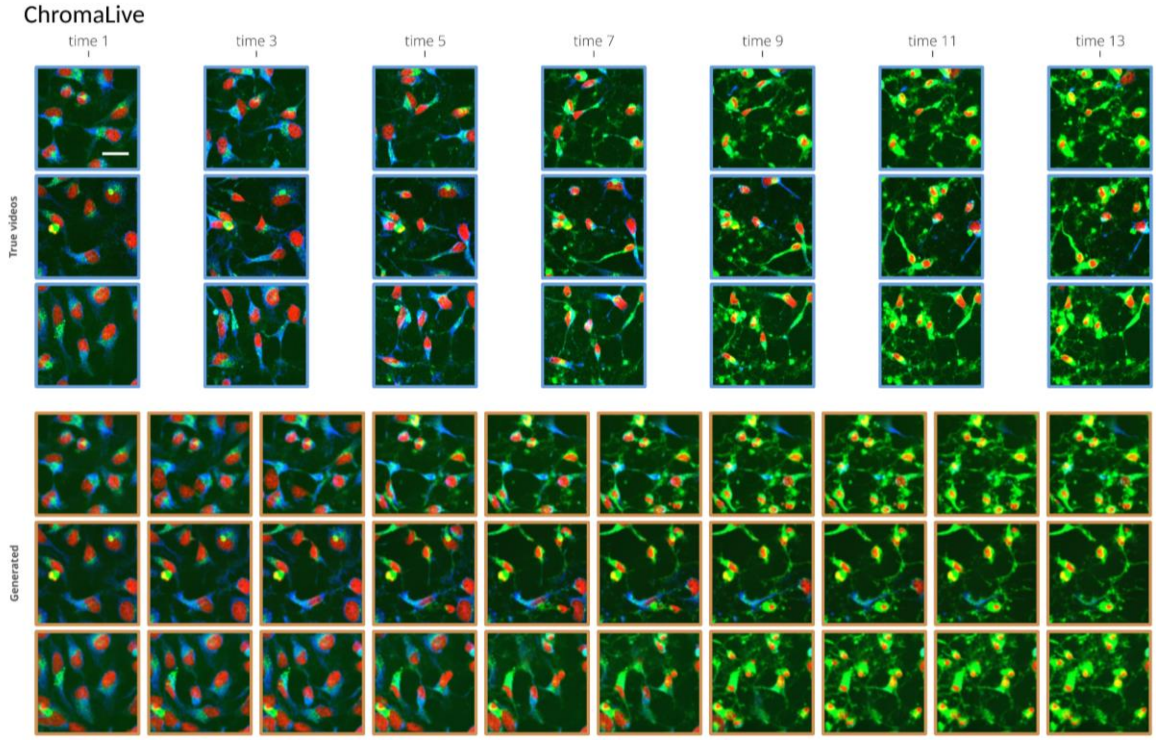


**Supplementary Figure6: True and generated videos for ChromaLive.** Columns: subsampled video time steps. Top 3 rows: 3 true videos. Bottom 3 rows: 3 generated videos from the same first image. The scale bar is 30 µm.

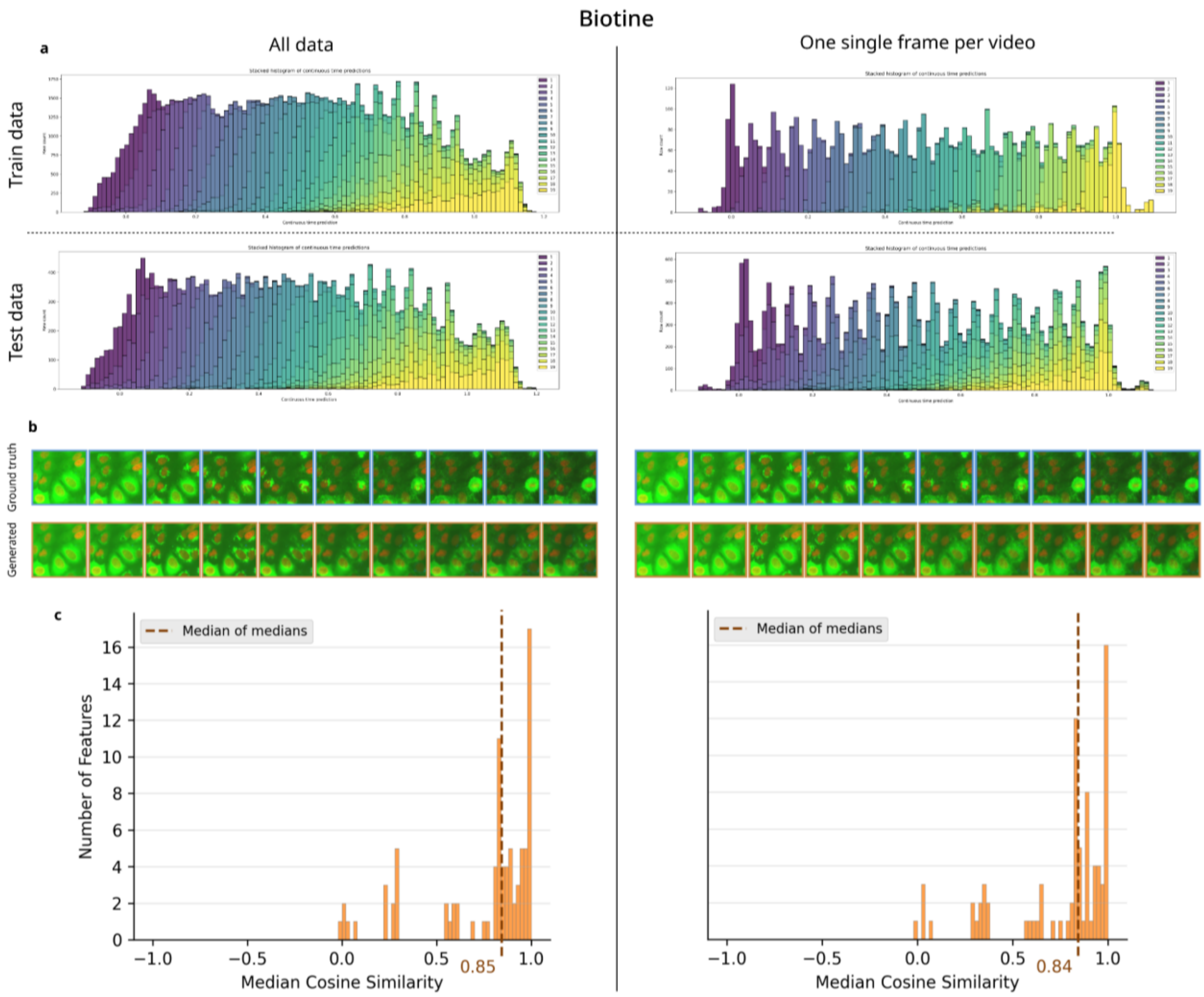


**Supplementary Figure7: Static2Dynamic is robust to time-unpaired data.** Left column: the full Biotine dataset. Right column: a subset of the Biotine dataset where a single frame was kept for each video (drastically lowering the amount of training data, but akin to a real dataset where sequences are not available). a - histograms of predicted pseudotimes for both train sets and for a common test set between the two experiments (including paired frames from full videos). b - examples of true video (common to both test sets), and the corresponding generated video. C - CellProfiler evaluations akin to **Figure 2**G (see **Supp. Fig. 11** for details).

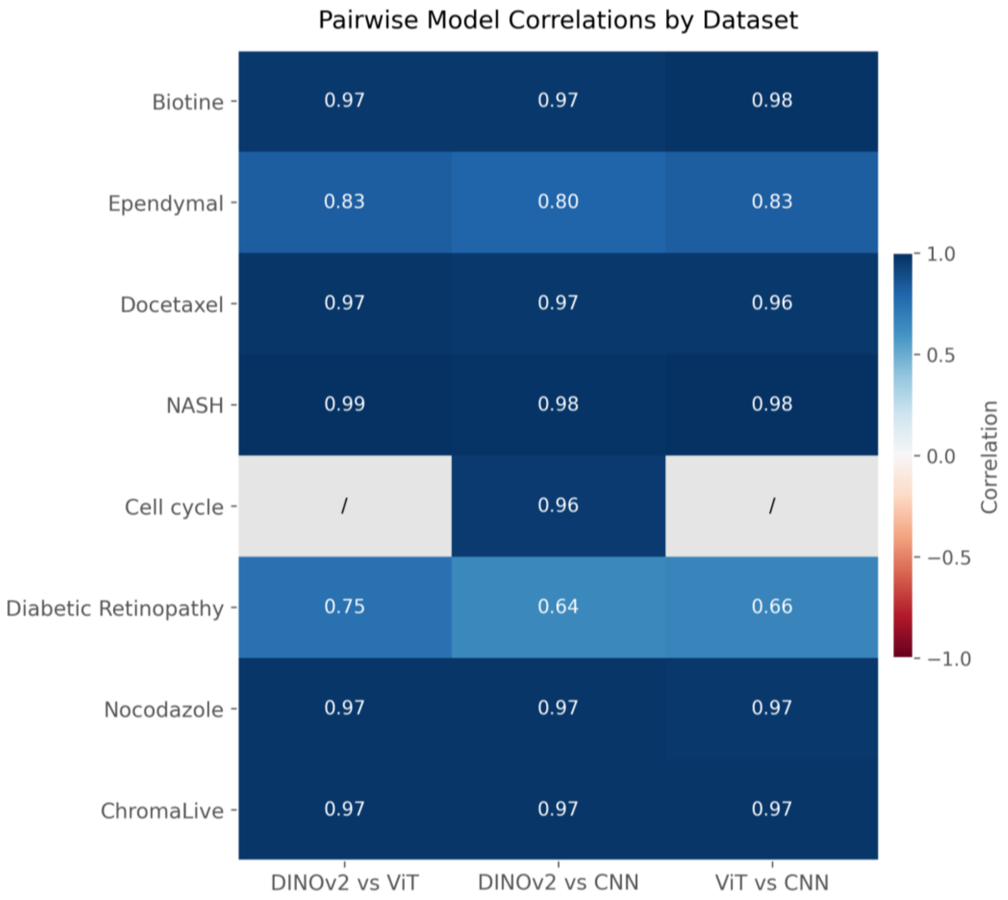


**Supplementary Figure8: Pseudotime estimation yields similar results across different image encoders.** The correlation between the predicted pseudotimes obtained using any two different image encoders is very high, especially on datasets where the predicted pseudotimes exhibit very high Kendall Tau scores w.r.t. ground truth labels. CNN is `timm/convnextv2_large.fcmae`, `VIT is google/vit-huge-patch14-224-in21k` and DINOV2 is `facebook/dinov2-with-registers-giant`.

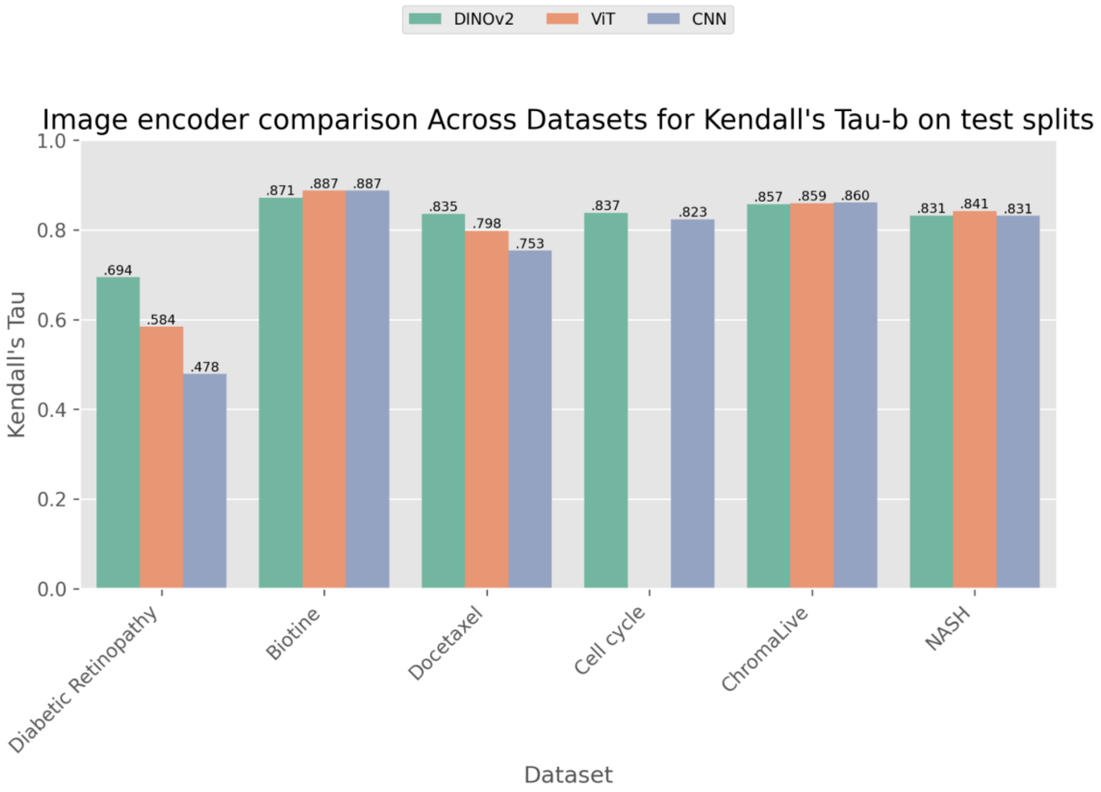


**Supplementary Figure9: Pseudotime estimation rank is robust to the image encoder.** Kendall Tau statistic between predicted pseudotimes and ground truth labels. While most encoders provide a pseudotime rank that aligns with ground truth, DINOv2 appears to be slightly better on some dataset. This is especially the case on Diabetic Retinopathy, maybe due to the fact it exhibits a high image variability. The Ependymal dataset is not included because the amount of data for which we have ground truth labels (to compute the Kendall Tau from ground truth here) is limited. The VIT model could not be used for the Cell Cycle dataset because its standard processing pipeline does not support single-channel images. CNN is `timm/convnextv2_large.fcmae`, `VIT is google/vit-huge-patch14-224-in21k` and DINOV2 is `facebook/dinov2-with-registers-giant`.

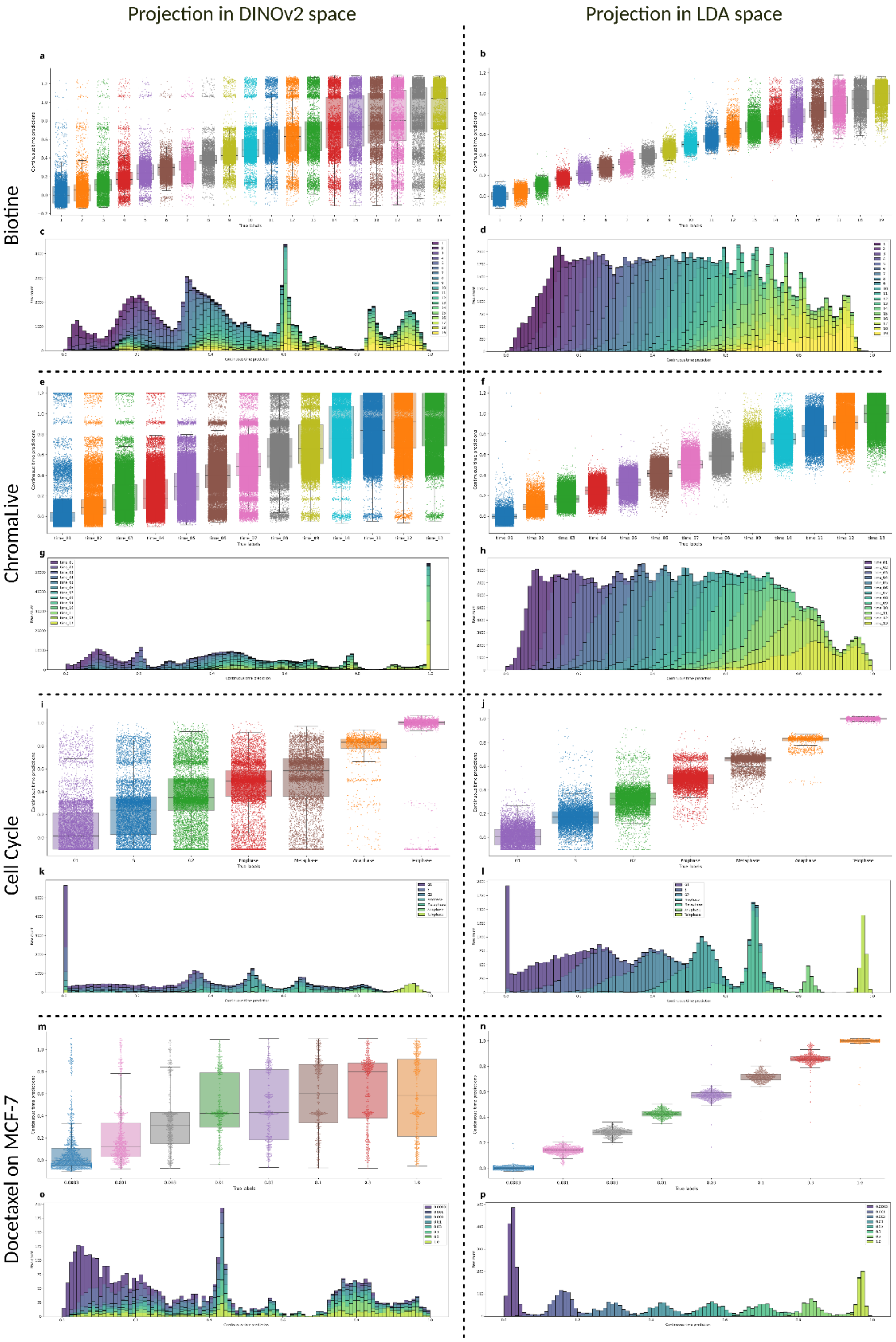


**Supplementary Figure10: The LDA transform is key to the first stage of Static2Dynamic.** Shown here for four datasets (in rows) are boxplots and histograms of the continuous pseudotime predictions obtained by Static2Dynamic on the right column, and by an ablation model where the projections onto the time spline were made directly in the DINO embedding space (instead of the LDA space) in the left column. The LDA transform provides a more discriminative subspace than the raw DINO embedding from which a robust continuous pseudotime can be extracted.

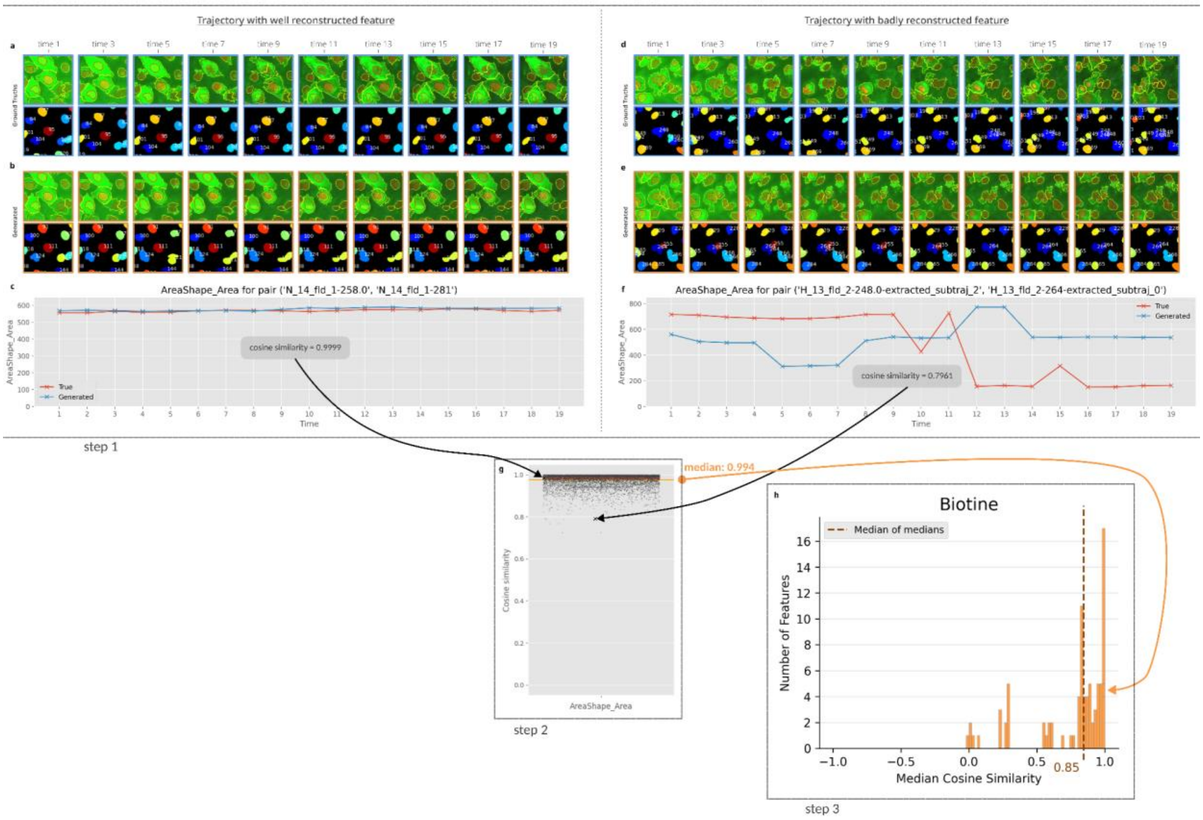


**Supplementary Figure11: Evaluation pipeline for the recovery of unseen dynamics based on Cellprofiler features.** In order to evaluate Static2Dynamic on its ability to recover the *unseen dynamics* of ground truth data, we generated videos matching real videos by starting from the same initial frame for both Biotine and ChromaLive datasets. We then segmented and tracked each cell along time, and computed CellProfiler (CP) (Stirling et al. 2021) features on these single-cell video cutouts (step 1). We then paired each true cell trajectory with its generated counterpart based on the initial position. We then compared the obtained CP time-indexed feature vectors of each true versus generated cell pair by computing the cosine similarity between their feature vector, for each feature (step 2). In order to not artificially favor the evaluation, we removed redundant highly correlated features as well as features that did not show significant variations along time in the first place (see **Supp. Fig. 23**). We then report the histogram of median cosine similarities across all cell pairs, for each feature (step 3).

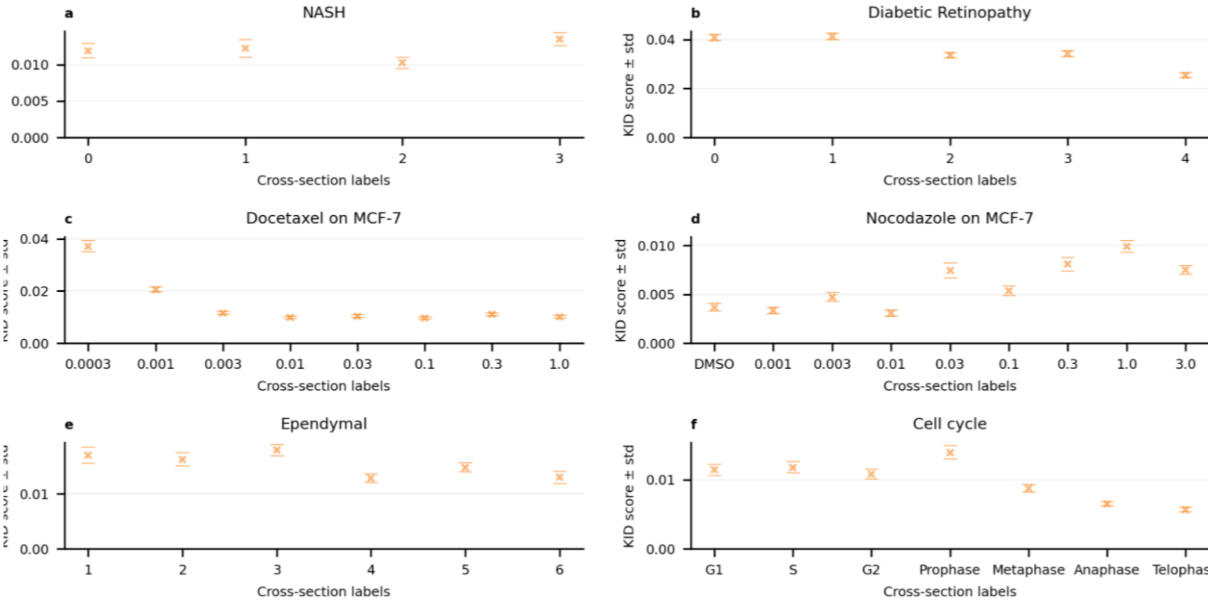


**Supplementary Figure12: KID scores for all datasets.** KID scores at each cross section label is computed the following way: 1) pseudotime are estimated for all real images available for the considered cross section, 2) as many images are generated from these pseudotime with a random seed, 3) KID is computed between these two datasets. Standard deviation is over 100 random subsets of InceptionV3 features.

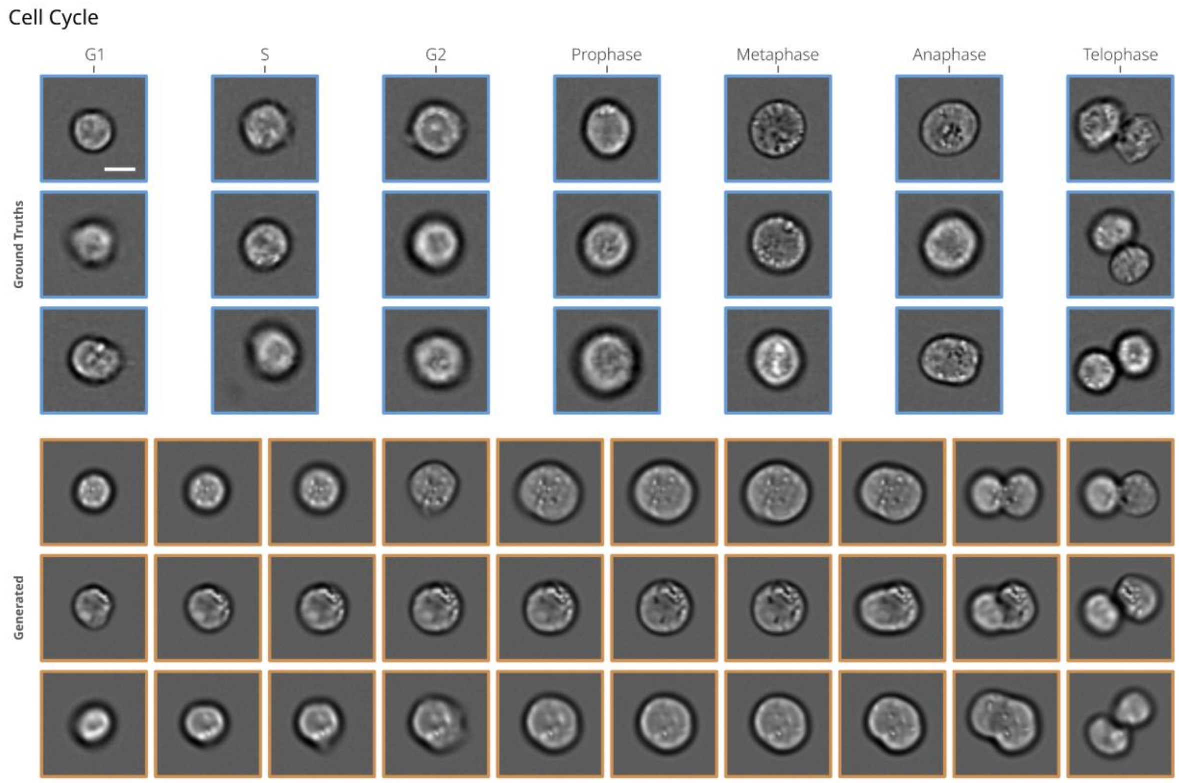


**Supplementary Figure13: Examples of true image samples and generated videos for the Cell Cycle dataset.** The scale bar is 10 µm.

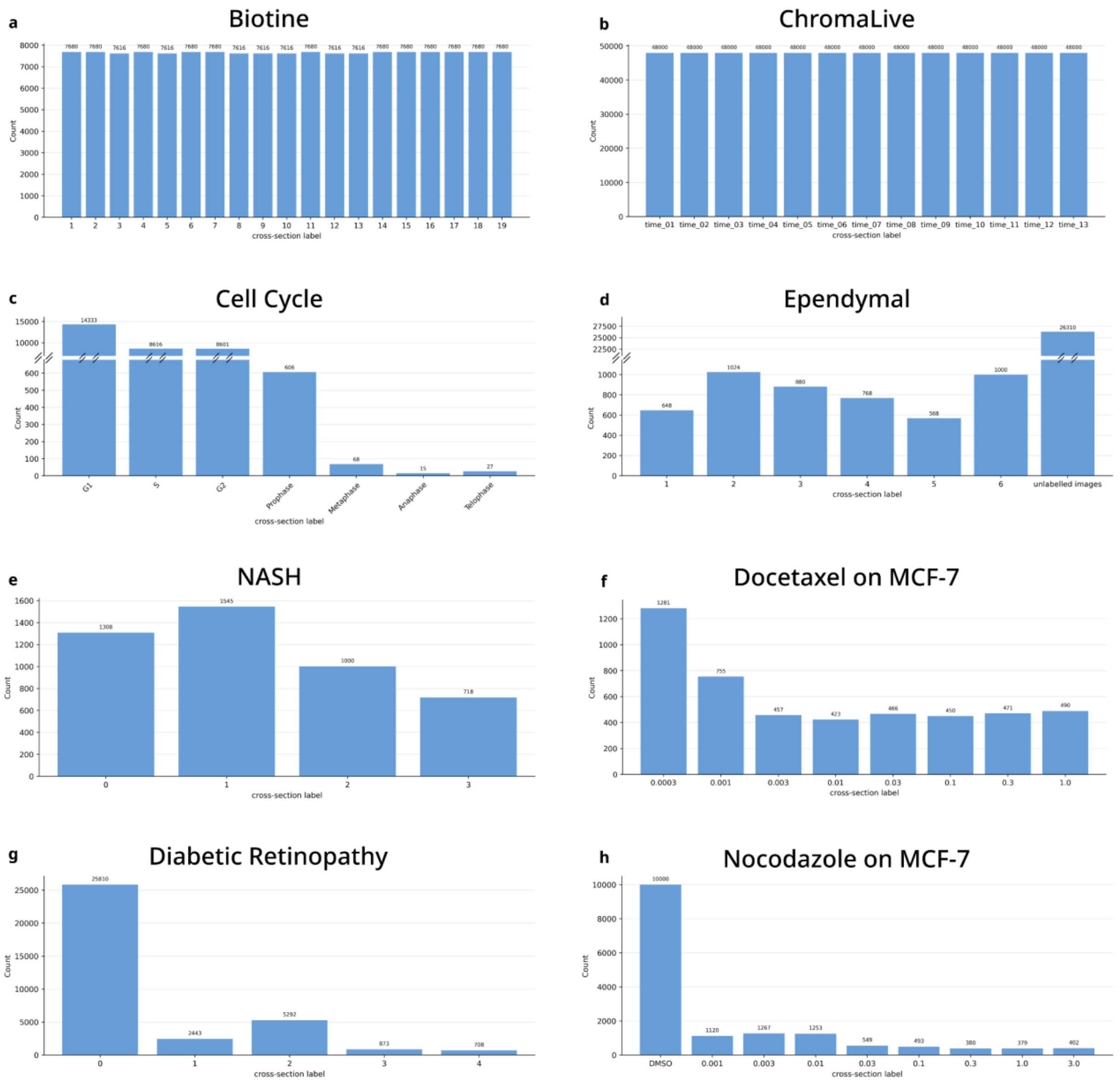


**Supplementary Figure14: Histograms of true image counts per cross-section label for all datasets.** These counts are before the 8x data augmentation (for the Biotine, NASH, Docetaxel and Nocodazole datasets) or continuous circle rotation augmentations (for Cell Cycle, Ependymal, and Diabetic retinopathy) is performed. For the Biotine dataset, one specific field of view (= 64 images) was removed at some specific times because it was blurry. For the ChromaLive dataset, the images used for training are crops taken at random positions and orientations over the full fields of view.

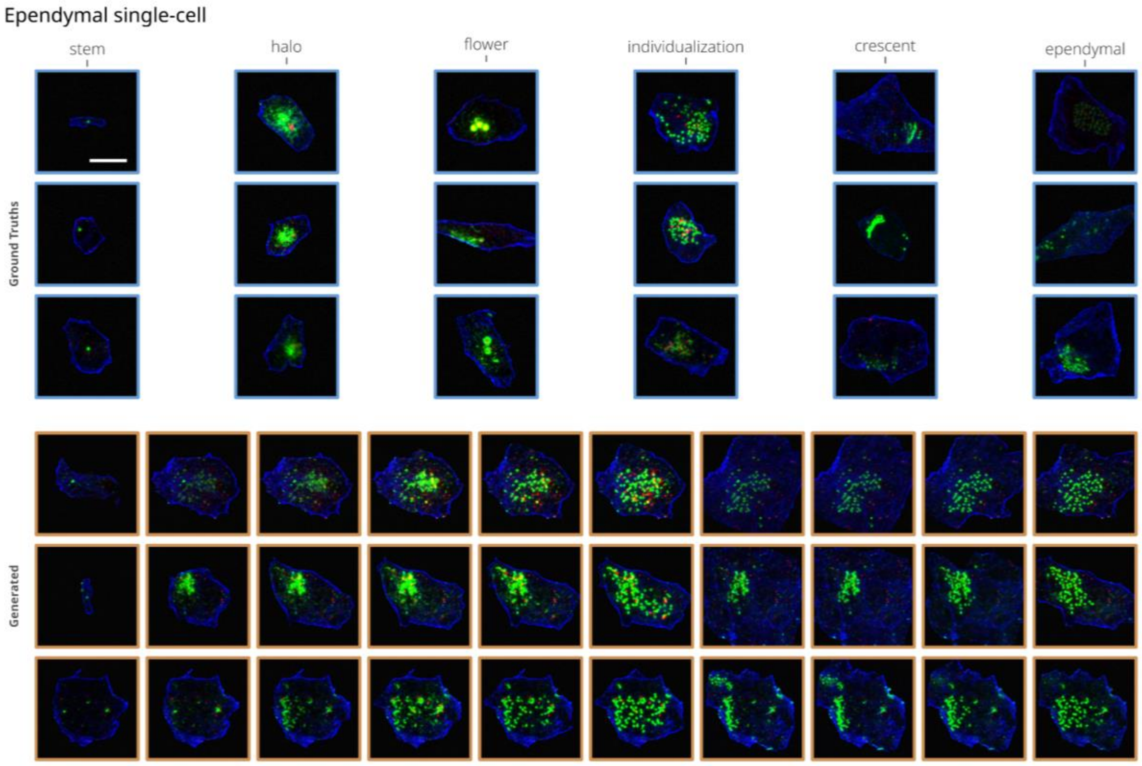


**Supplementary Figure15: Examples of true image samples and generated videos for the Ependymal dataset.** The scale bar is 20 µm.

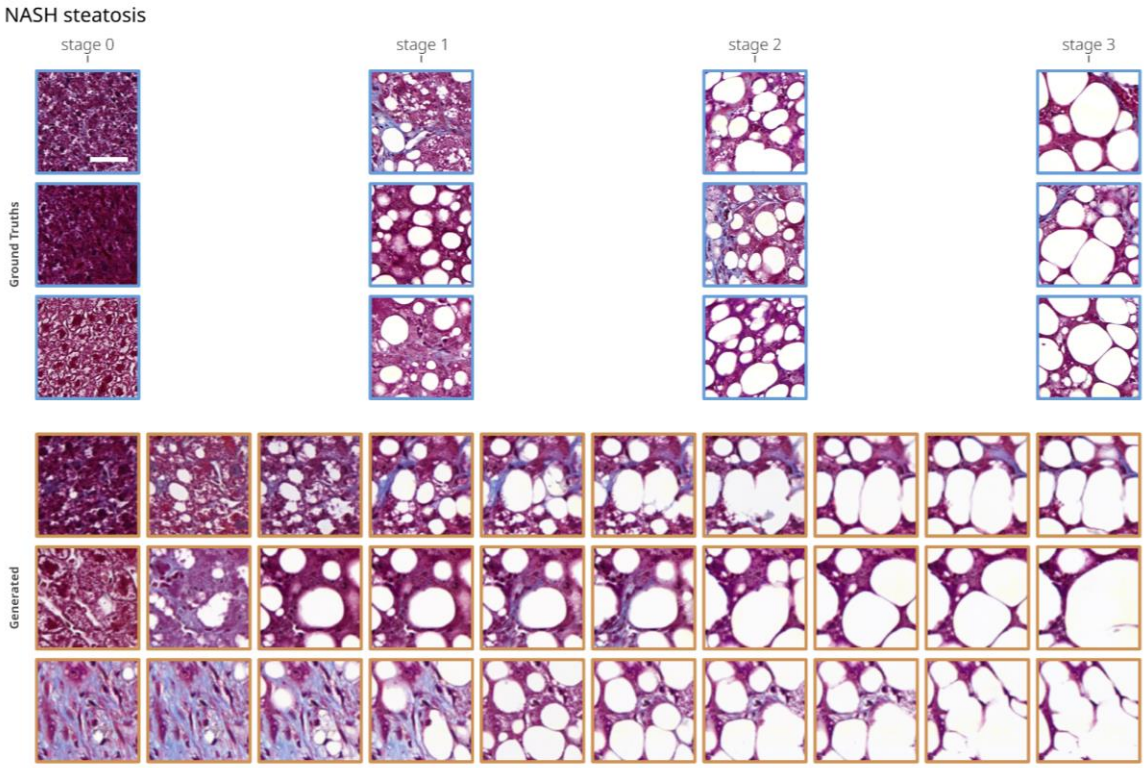


**Supplementary Figure16: Examples of true image samples and generated videos for the NASH dataset.** The scale bar is 50 µm.

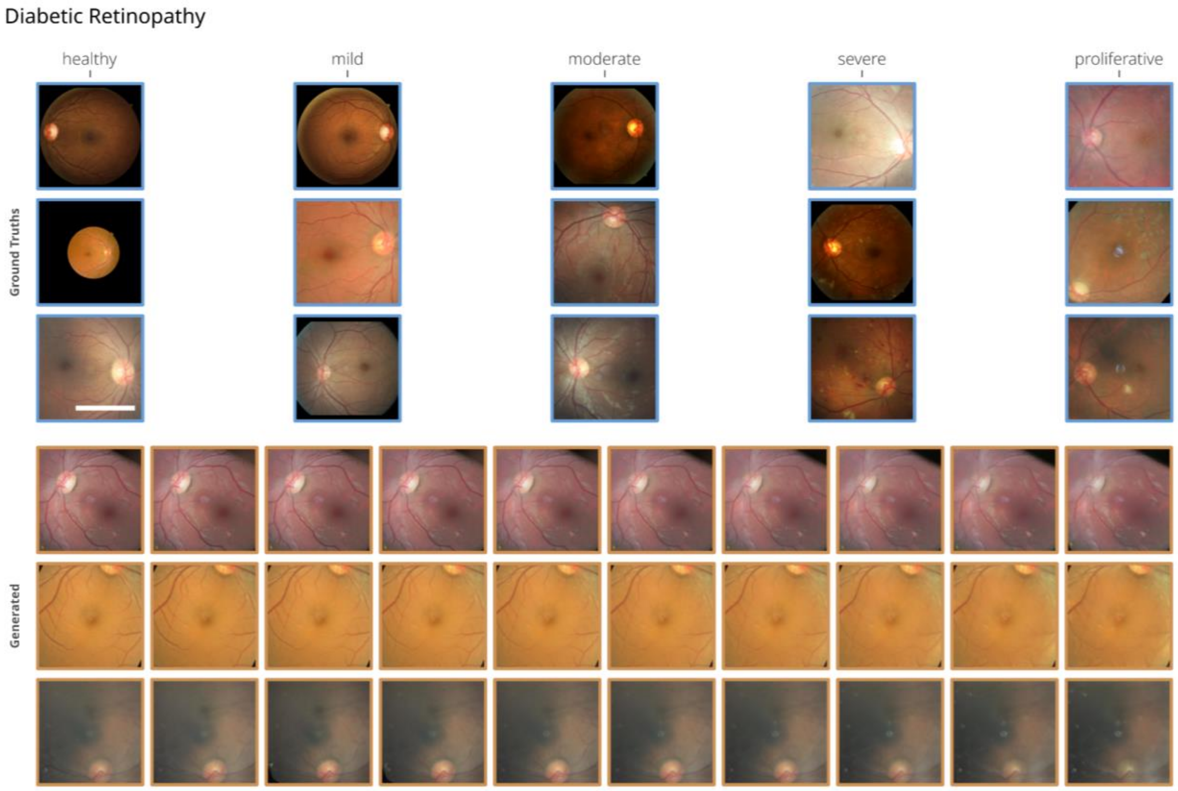


**Supplementary Figure17: Examples of true image samples and generated videos for the Diabetic Retinopathy dataset.** This dataset exhibits highly heterogeneous pixel scales; for reference, the scale bar is around 5 mm for the first image of the third row.

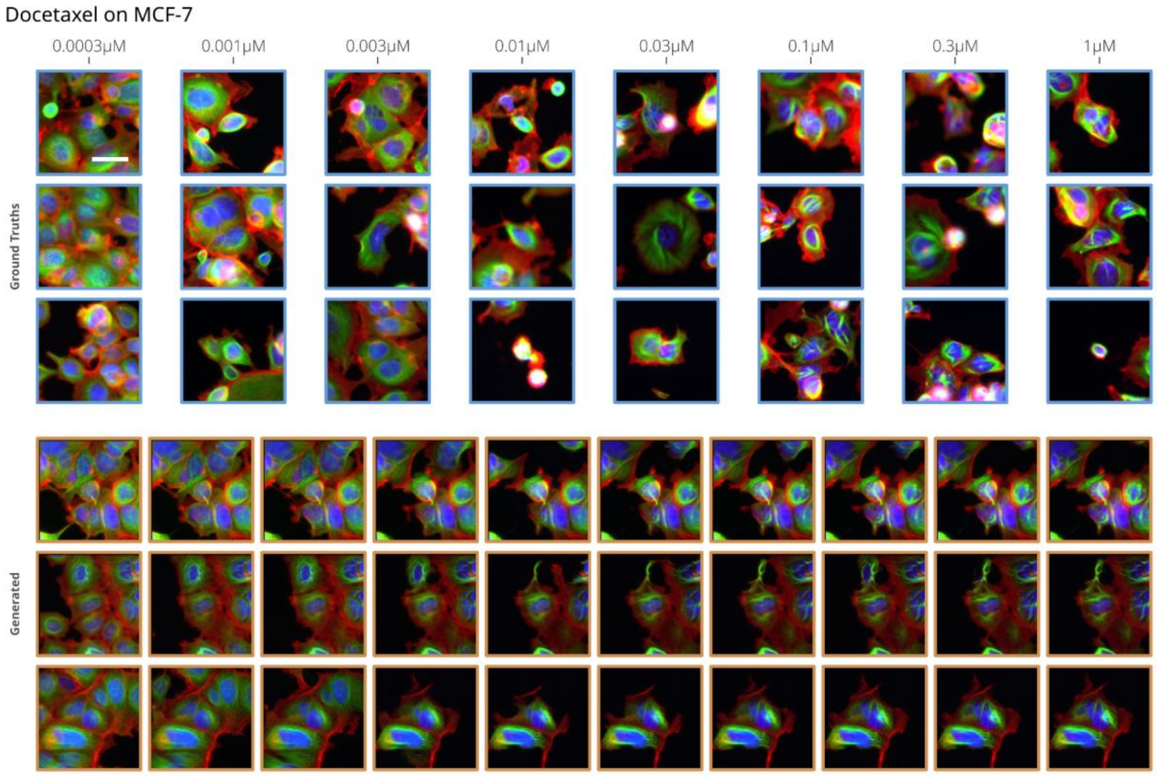

**Supplementary Figure18: Examples of true image samples and generated videos for the Docetaxel on MCF-7 dataset.** The scale bar is 30 µm.

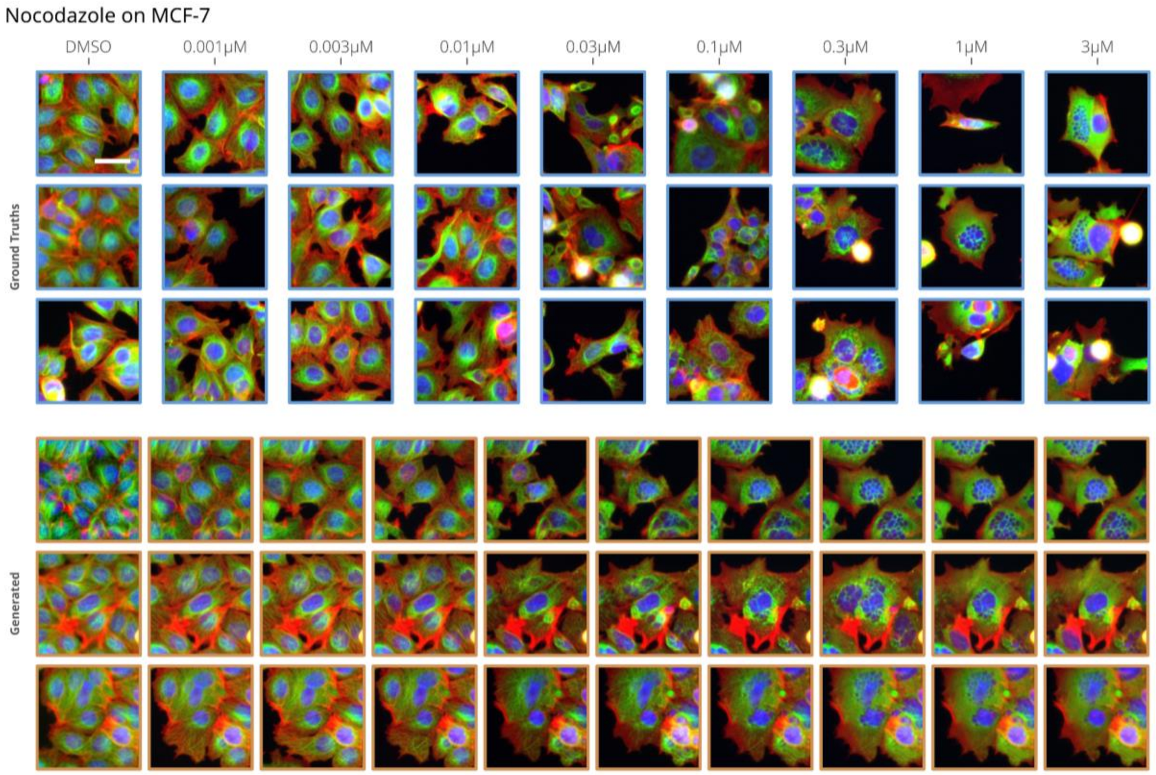


**Supplementary Figure19: Examples of true image samples and generated videos for the Nocodazole on MCF-7 dataset.** The scale bar is 30 µm

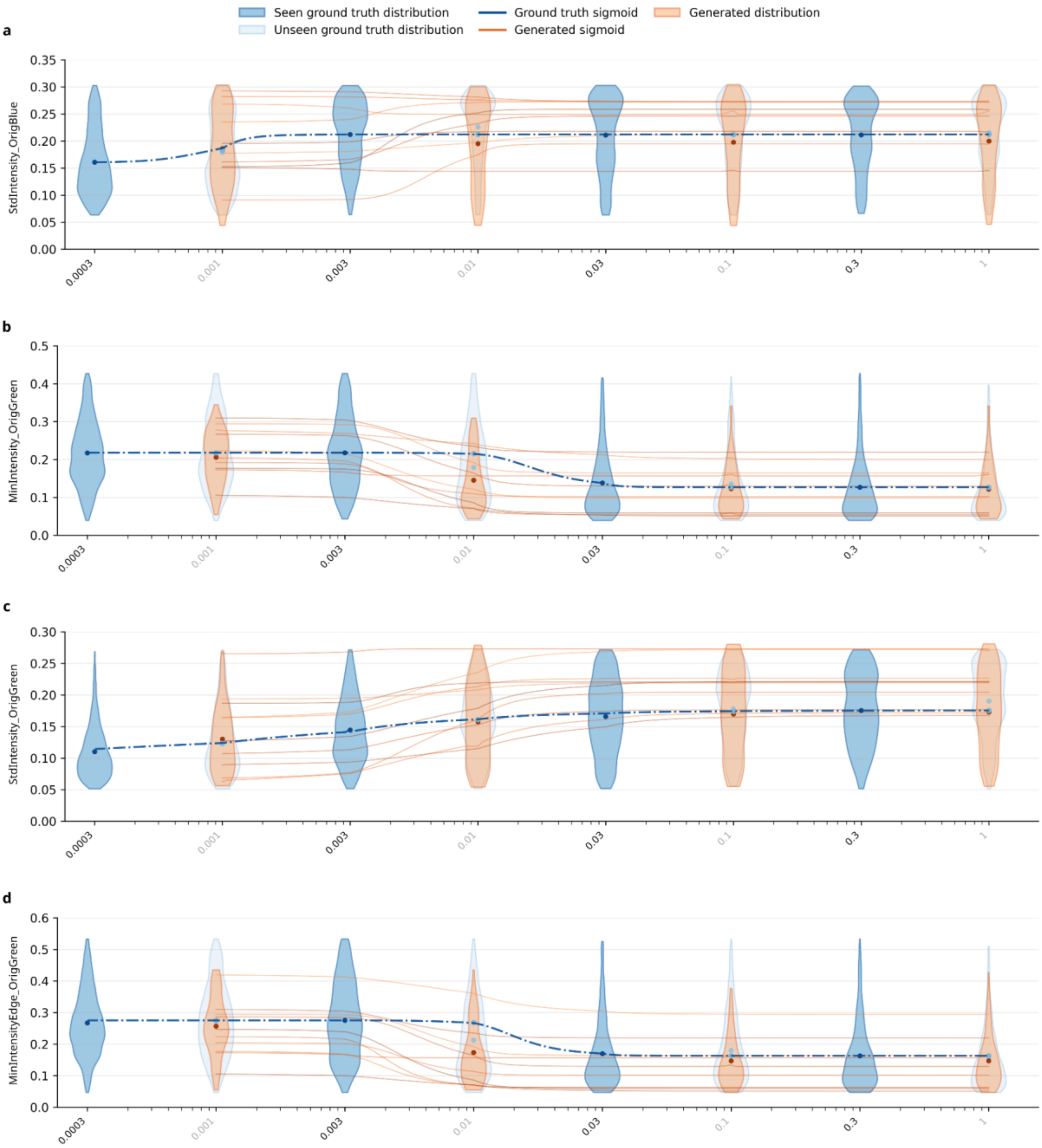


**Supplementary Figure20: Examples of four continuous single cell feature dose responses on Docetaxel on the MCF-7 dataset.** In blue, distribution of the feature is computed on real data that is seen by the model, in orange, data unseen by the model. Static2Dynamic makes possible the computation of a continuous dose response for any quantitative feature on any single cell (orange lines) which may differ from an arbitrary sigmoid model fit on the average values of 4 doses (dotted blue lines) as usually performed.

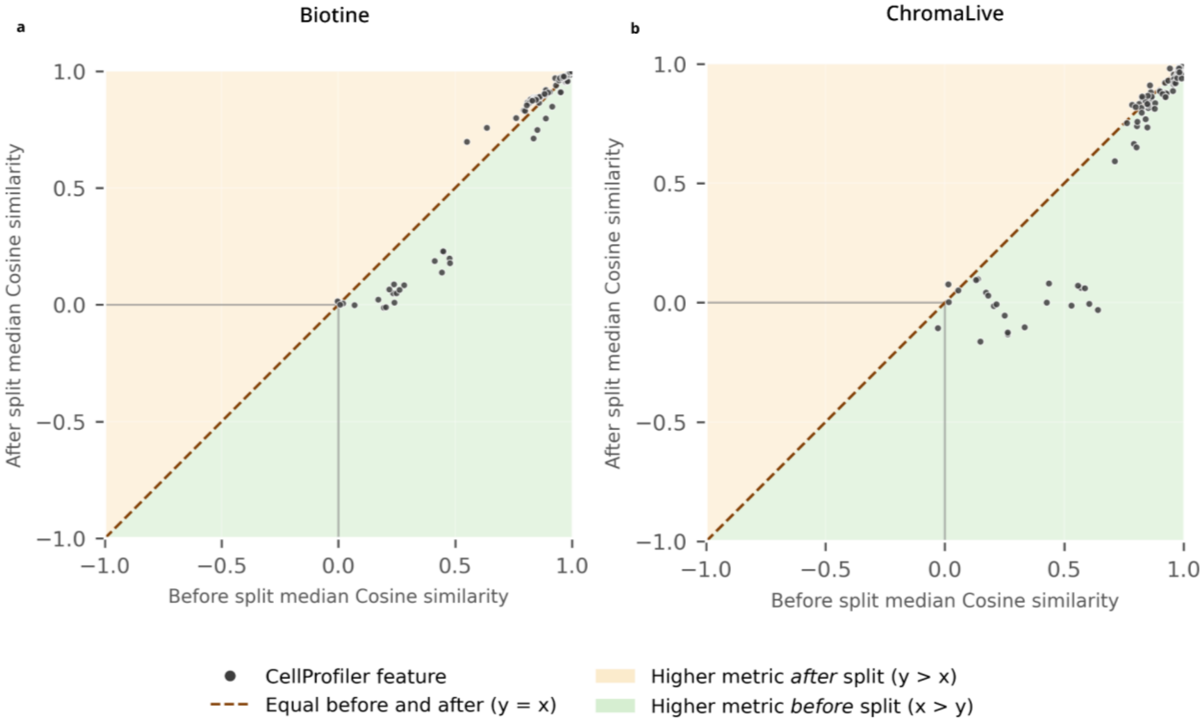


**Supplementary Figure21: Feature cosine similarity is degraded after cell division.** Here we focused the analysis on pairs of tracks where a cell division event was reported on the real data by the CellProfiler tracking. We then computed the median cosine similarities on the trajectory parts *before* versus *after* the split time, for all features. These results underline the fact that Static2Dynamic doesn't properly recapitulate the features of the daughter cells after splitting because the movie generation didn't produce the cell division.

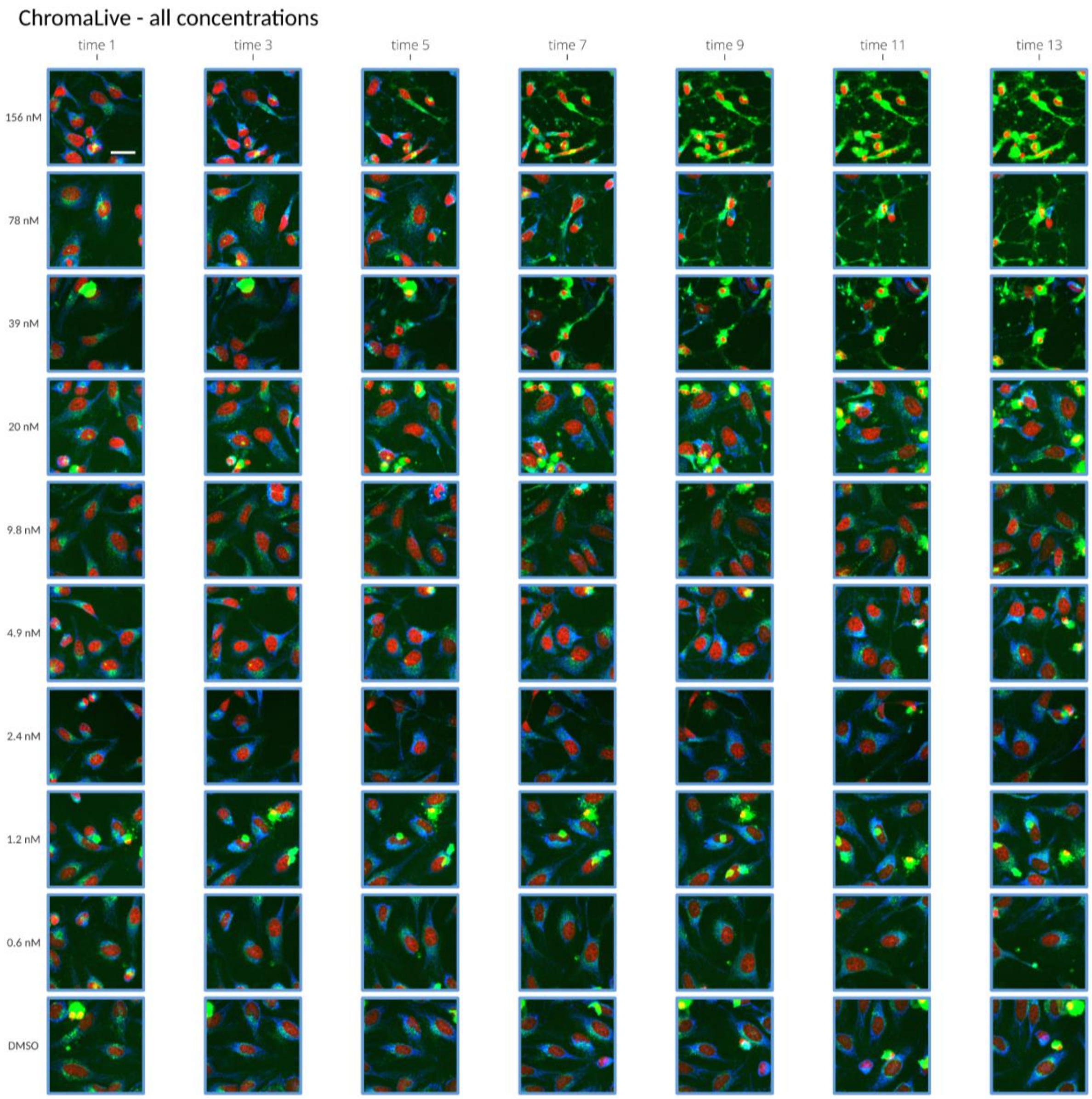


**Supplementary Figure22: The full ChromaLive assay at all concentrations.** Rows: concentration of compound, by decreasing order. Columns: video times. The low concentrations induce an extremely subtle phenotype, contrary to the highest ones. Static2Dynamic is trained on the 2 highest concentrations together. The scale bar is 30 µm.

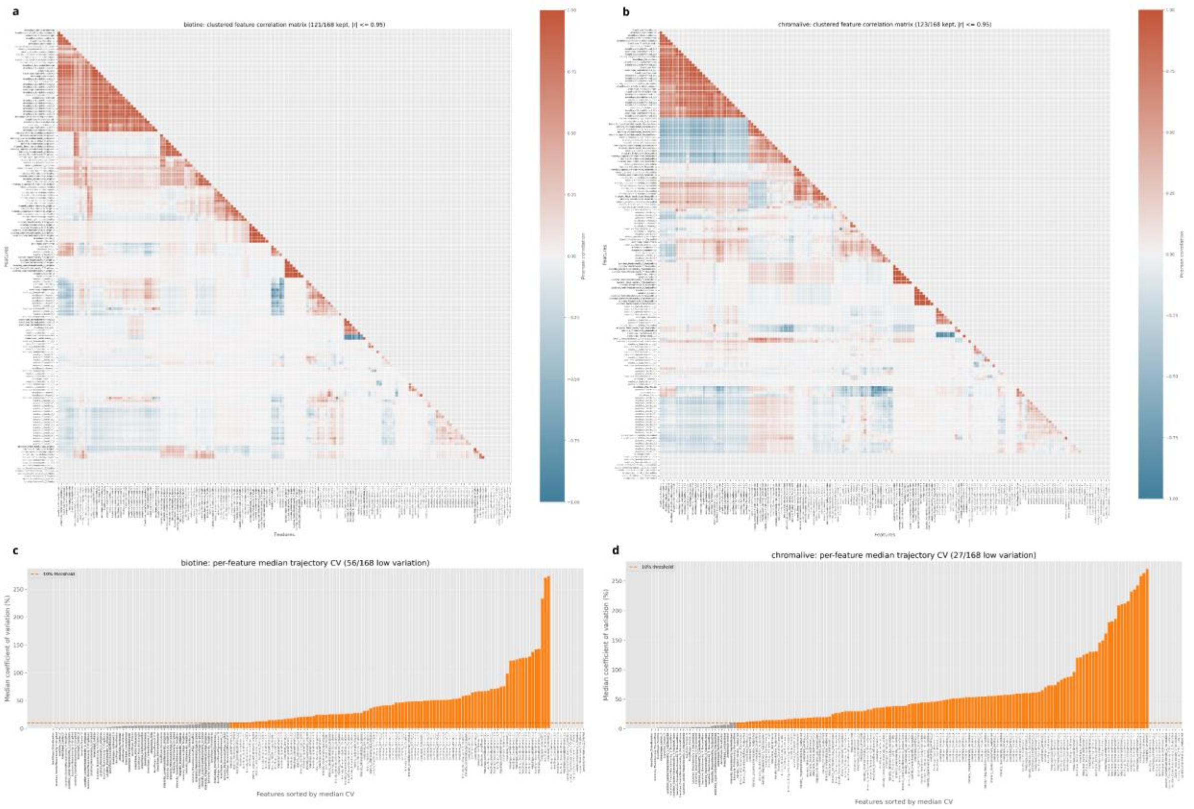


**Supplementary Figure23: CellProfiler features cross-correlations and Coefficients of Variation (CV).** Bold features were removed from the set of considered features for the CellProfiler analysis.
